# Near telomere-to-telomere genome assemblies of two *Chlorella* species unveil the composition and evolution of centromeres in green algae

**DOI:** 10.1101/2024.02.27.582245

**Authors:** Bo Wang, Yanyan Jia, Ningxin Dang, Jie Yu, Stephen J. Bush, Shenghan Gao, Wenxi He, Sirui Wang, Hongtao Guo, Xiaofei Yang, Weimin Ma, Kai Ye

## Abstract

**Background:** Centromeres play a crucial and conserved role in cell division, although their composition and evolutionary history in green algae, the evolutionary ancestors of land plants, remains largely unknown.

**Results:** We constructed near telomere-to-telomere (T2T) assemblies for two Trebouxiophyceae species, *Chlorella sorokiniana* NS4-2 and *Chlorella pyrenoidosa* DBH, with chromosome numbers of 12 and 13, and genome sizes of 58.11 Mb and 53.41 Mb, respectively. We identified and validated their centromere sequences using CENH3 ChIP-seq and found that, similar to humans and higher plants, the centromeric CENH3 signals of green algae display a pattern of hypomethylation. Interestingly, the centromeres of both species largely comprised transposable elements, although they differed significantly in their composition. Species within the *Chlorella* genus display a more diverse centromere composition, with major constituents including members of the LTR/Copia, LINE/L1, and LINE/RTEX families. This is in contrast to green algae including *Chlamydomonas reinhardtii*, *Coccomyxa subellipsoidea*, and *Chromochloris zofingiensis*, in which centromere composition instead has a pronounced single-element composition. Moreover, we observed significant differences in the composition and structure of centromeres among chromosomes with strong collinearity within the *Chlorella* genus, suggesting that centromeric sequence evolves more rapidly than sequence in non-centromeric regions.

**Conclusions:** This study not only provides high-quality genome data for comparative genomics of green algae but gives insight into the composition and evolutionary history of centromeres in early plants, laying an important foundation for further research on their evolution.

## Background

Centromeres play a critical role in ensuring the accurate segregation of chromosomes during both mitosis and meiosis [1], being specialized chromosomal regions associated with chromatid cohesion and correct separation during cell division. Centromeres serve as the assembly and attachment points for the kinetochore, a large protein complex that facilitates the binding of spindle microtubules to chromatids, enabling their movement towards the poles [2]. The ‘centromere paradox’ refers to the apparent contradiction between this essential role in chromosome segregation, which is conserved across all eukaryotes, and the significant variability in their organization and DNA composition, even within closely related species [3, 4]. This variability highlights the (‘paradoxically’) rapid evolution of centromeres.

The high prevalence of satellite repeats within centromere sequences also presents a considerable genome assembly challenge, frequently leading to the substantial loss of sequence information in this region [5]. This obstacle poses a significant impediment to the comprehensive investigation of centromere functionality. However, with advances in sequencing technology, in particular the PacBio high-fidelity (HiFi) and Oxford Nanopore Technologies (ONT) ultra-long methods, in conjunction with advances in assembly algorithms [6, 7], significant progress has been made towards the complete assembly of centromeres in many model species, including but not limited to human [8], *Arabidopsis thaliana* [9, 10] rice [11], and maize [12]. Notably, both human and higher plant centromeres primarily comprise numerous satellite sequences, organized into higher-order repeat arrays [5, 13]. It has previously been proposed that plant centromeric satellite sequences may have arisen from transposable element (TE) insertions specifically targeting the centromere [14, 15]. Consistent with this, the centromeres of the oldest known domesticated wheat, *Triticum monococcum*, have recently been found to predominantly feature transposable elements [16], as have those of the moss *Physcomitrium patens* and the model green alga *Chlamydomonas reinhardtii* [17, 18]. Whether centromeres in other green algae, more distantly related to *Chlamydomonas reinhardtii*, are also composed of TEs, and if so, whether the constituent elements are similar, as well as the evolutionary transitions between them, remains unclear.

In this study, we sequenced and assembled two *Chlorella* species to near-T2T level, characterising and experimentally validating their centromeres, and comparing them to those of *Chlamydomonas reinhardtii, Coccomyxa subellipsoidea*, and *Chromochloris zofingiensis*. We found that the centromeres of the green algae species selected in this study extensively comprise TEs, albeit with significant differences in composition. *Chlamydomonas reinhardtii* and *Coccomyxa subellipsoidea* predominantly feature elements from the LINE/L1 family, while *Chromochloris zofingiensis* primarily comprises elements from the LTR/Copia family. In contrast, species within the *Chlorella* genus exhibit a more diverse centromere composition, the major constituents of which were members of the LTR/Copia, LINE/L1, and LINE/RTEX families. Overall, our results give insight into the centromere evolution of green algae, which may be generalised to early plants, as well as providing high-quality genomic resources to facilitate future comparative analyses.

## Results

### Near telomere-to-telomere assemblies of two *Chlorella* species

To investigate centromere evolution within the context of green algae, we assembled the genomes of two *Chlorella* species, *Chlorella sorokiniana* NS4-2 and *Chlorella pyrenoidosa* DBH (basionym: *Auxenochlorella pyrenoidosa*), both of which belong to the class Trebouxiophyceae (phylogenetically distant from the model green alga *Chlamydomonas reinhardtii*, with a divergence time of 827 MYA; http://timetree.org/) (**Supplementary Fig. S1**). The two species are morphologically similar, although vary in their chromosome numbers, with *Chlorella sorokiniana* NS4-2 having 12 chromosomes and *Chlorella pyrenoidosa* DBH having 13 (**Fig. 1A**, **Fig. 1B, Supplementary Fig. S2A and Supplementary Fig. S2B**). The genomes of *Chlorella sorokiniana* NS4-2 and *Chlorella pyrenoidosa* DBH were sequenced using HiFi reads at high coverage depths of 438 and 308-fold respectively (**Supplementary Table S1**). For *Chlorella sorokiniana*, we assembled the HiFi reads (**Supplementary Fig. S3A**) to produce a set of 16 contigs. These contigs had a minimum length of 141,360 bp and were selected based on the criterion of length exceeding 100 Kb. Following this, we generated a final set of 13 contigs by excluding the genome of the putative alga-benefitting commensal bacterium *Arthrobacter* sp. [19] as well as fragments of rDNA and plastid DNA (21 Kb∼107 Kb). Among these contigs, 11 contained the plant-type telomeric repeat unit (CCCTAAA/TTTAGGG) at both ends, signifying their completeness. Additionally, two contigs were found to have rDNA units at one end, as observed through collinearity analysis in comparison with the closely related species *Chlorella vulgaris* which was scaffolded by optical mapping [20] (**Supplementary Fig. S4A**). To consolidate the rDNA ends originating from the same chromosome, we merged the two contigs and inserted 100 Ns as spacers. As a result of this process, we produced a near-completely T2T 12-chromosome assembly of *Chlorella sorokiniana* NS4-2, the only gaps in which were in the rDNA region of chromosome 1. This conclusion was further supported by karyotyping results (**Fig. 1B**) and Hi-C contact maps (**Fig. 1C**).

**Fig. 1.**
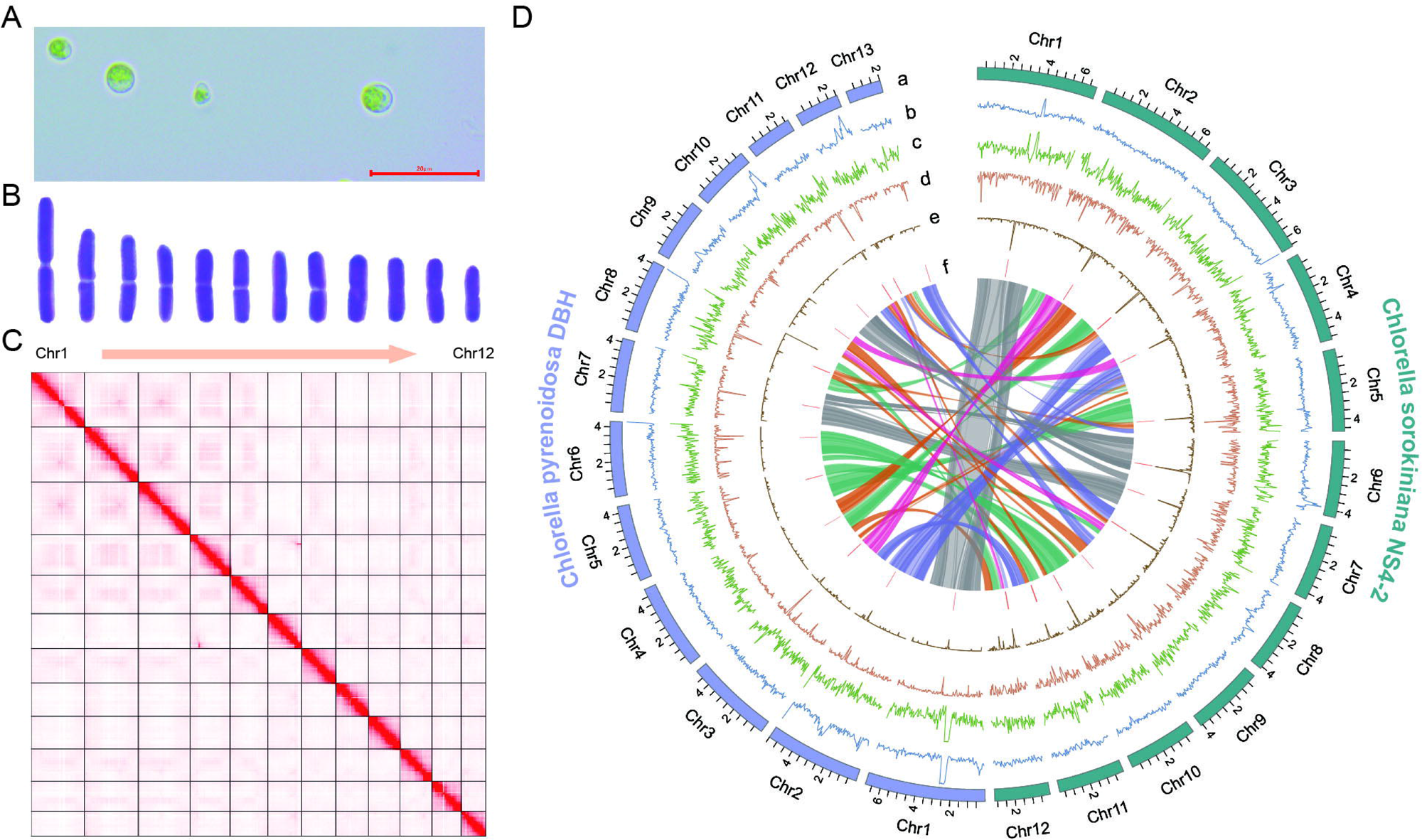
T2T genome assemblies of *Chlorella sorokiniana* NS4-2 and *Chlorella pyrenoidosa* DBH. (A) Micrograph of *C. sorokiniana* NS4-2, with bar length = 20 μm; **(B)** Karyotype diagram indicating that *C. sorokiniana* NS4-2 has 12 chromosomes; **(C)** Hi-C contact map of the *C. sorokiniana* NS4-2 T2T assembly; **(D)** Circular plot showing the principal genomic features of both the *C. sorokiniana* NS4-2 and *C. pyrenoidosa* DBH assemblies: karyotype distribution (track a); GC density (track b); gene density (track c); LINE element density (track d); LTR element density (track e); putative centromeric loci (track f). Densities were calculated in 50 Kb windows. The innermost of the circular plots represents the synteny relationship between *C. sorokiniana* NS4-2 and *C. pyrenoidosa* DBH.

For *Chlorella pyrenoidosa* DBH, we initially assembled the HiFi reads using hifiasm [6] (**Supplementary Fig. S3B**), which generated 38 contigs, each exceeding 100 Kb in length. However, upon closer examination, we discovered that only three of these contigs contained the telomeric repeat unit at both ends. This finding suggests that achieving a T2T-level assembly for *Chlorella pyrenoidosa* DBH is more challenging compared to *Chlorella sorokiniana* NS4-2 (**Supplementary Fig. S3A**). To overcome this challenge, we generated ONT ultra-long reads (with a read length N50 of 59 Kb) at a high coverage depth of 368-fold (**Supplementary Table S1**), assembling them using NextDenovo [21] to generate a set of 14 contigs. Of these 14 contigs, 10 exhibited the presence of telomeric repeat units at both ends, indicating the complete resolution of the telomere regions. However, two contigs only had rDNA units at one end and two only had the telomeric repeat unit at one end, in both cases suggesting an unresolved telomeric region at the other. To address this issue and achieve a more comprehensive assembly, we utilized the HiFi contigs to replace the non-resolved ONT contigs (**Supplementary Fig. S3B**). By incorporating these updated HiFi contigs, we successfully achieved a gap-free assembly of 14 contigs. This updated assembly provides a more complete representation of the genome, with improved resolution and continuity. In a similar vein, for *Chlorella pyrenoidosa* DBH, collinearity analysis also revealed two contigs with rDNA units potentially originating from the same chromosome (**Supplementary Fig. S4B**). Following a similar approach to *Chlorella sorokiniana* NS4-2, the rDNA ends were merged, and 100 Ns were introduced as spacers. This led to the generation of a virtually T2T assembly with 13 chromosomes, again supported by karyotyping results (**Supplementary Fig. S2B**) and Hi-C contact maps (**Supplementary Fig. S2C**).

Genome completeness was evaluated using Benchmarking Universal Single-Copy Orthologs (BUSCO), with *Chlorella sorokiniana* NS4-2 scoring 98.3% and *Chlorella pyrenoidosa* DBH 99.0%. The assembly of these two genomes therefore represents a significant improvement compared to existing genomes for these species (**Table 1**). *Chlorella sorokiniana* NS4-2 and *Chlorella pyrenoidosa* DBH were predicted to have 12,691 and 11,327 protein coding genes, respectively, and to exhibit a significant collinearity relationship, accompanied by chromosomal rearrangements (**Fig. 1D**). Finally, all HiFi reads greater than 15 Kb were selected and used to perform mitochondrial and chloroplast (i.e. circular) genome assembly using Unicycler v0.5.0 with default parameters [22]. The chloroplast genome sizes of *Chlorella sorokiniana* NS4-2 and *Chlorella pyrenoidosa* DBH are 109,738 bp and 107,436 bp, respectively; while the mitochondrial genome sizes are 53,629 bp and 69,538 bp, respectively (GenBank accession: OP311589-OP311592).

**Table 1.** Genomic features of the *Chlorella sorokiniana* NS4-2 and *Chlorella pyrenoidosa* DBH assemblies.

| Genome features | NS4-2 | UTEX1230<sup>a</sup> | DBH | FACHB-9<sup>b</sup> |
| --- | --- | --- | --- | --- |
| Nuclear genome size (Mb) | 58.11 | 58.36 | 53.41 | 56.77 |
| No. of chromosomes | 12 | NA | 13 | NA |
| No. of contigs | 13 | 18 | 14 | 8193 |
| GC content (%) | 63.95 | 63.95 | 66.94 | 66.71 |
| Contigs N50 (bp) | 4,352,225 | 3,818,101 | 4,302,578 | 9757 |
| Scaffolds N50 (bp) | 4,714,146 | 4,091,730 | 4,383,467 | 1,392,758 |
| No. of protein-coding genes | 12,691 | 12,871 | 11,327 | 10,283 |
| Repeat content (%) | 8.84 | 8.89 | 8.91 | 5.46 |

<sup>a</sup> UTEX1230 is a strain of *Chlorella sorokiniana* [36].
<sup>b</sup> FACHB-9 is a strain of *Chlorella pyrenoidosa* [37].

### Putative centromere identification and CENH3 ChIP-seq validation

Recent advances in sequencing technologies have shed light on the crucial role of TEs in influencing the characteristics of centromeric sequences among green algal species [17, 23]. Notably, studies of *Chlamydomonas reinhardtii* CC-4532 (v. 6.1) [24] and *Coccomyxa subellipsoidea* C-169 (v. 3.0) [25] have demonstrated the prevalence of LINE/L1 elements within their genomes, as well as LTR/Copia elements in *Chromochloris zofingiensis* [26], further emphasizing the impact of TEs in shaping their organism’s centromeric regions. These TEs are strikingly abundant in the centromeres, forming a unique and discernible structure when performing dot plots for self-alignment analysis (**Supplementary Fig. S5**). Notably, within this structure, specific clusters stand out, such as the *Zepp*-like cluster in *Chlamydomonas reinhardtii* and the *Zepp* cluster in *Coccomyxa subellipsoidea* (**Supplementary Fig. S5**). Leveraging this distinctive characteristic, we conducted dot plot analyses on a per-chromosome basis to pinpoint the potential centromeric regions within the genomes of *Chlorella sorokiniana* NS4-2 (**Supplementary Fig. S6**) and *Chlorella pyrenoidosa* DBH (**Supplementary Fig. S7**). In the case of *Chlorella sorokiniana* NS4-2, we successfully validated putative centromeres for all 12 chromosomes using VerityMap [27] (**Supplementary Fig. S8**), with lengths ranging from 28,465 bp to 104,090 bp (average length: 60 Kb) (**Supplementary Table S2**). Subsequently, we employed centromeric Histone H3 chromatin immunoprecipitation followed by sequencing technology (CENH3 ChIP-seq) to validate the predicted potential centromere regions and found them to be consistent with experimental verification (**Fig. 2 and Supplementary Fig. S9**). We then proceeded to use the dot plot method (**Supplementary Fig. S10**) for potential centromere prediction in *Chlorella pyrenoidosa* DBH species. For *Chlorella pyrenoidosa* DBH, we validated putative centromeres for all 13 chromosomes using VerityMap (**Supplementary Fig. S11**), with lengths ranging from 19,117 bp to 65,960 bp (average length: 39 Kb) (**Supplementary Table S3**). Insights from circular plots indicate that the two species exhibit a strong degree of collinearity and that the putative centromeres of *Chlorella sorokiniana* NS4-2 and *Chlorella pyrenoidosa* DBH are composed of TE elements (**Fig. 1D**). This was especially apparent for *Chlorella sorokiniana* NS4-2, in which the centromere regions were significantly enriched with LTR/Copia elements (**Fig. 2 and Fig. 1D**).

**Fig. 2.**
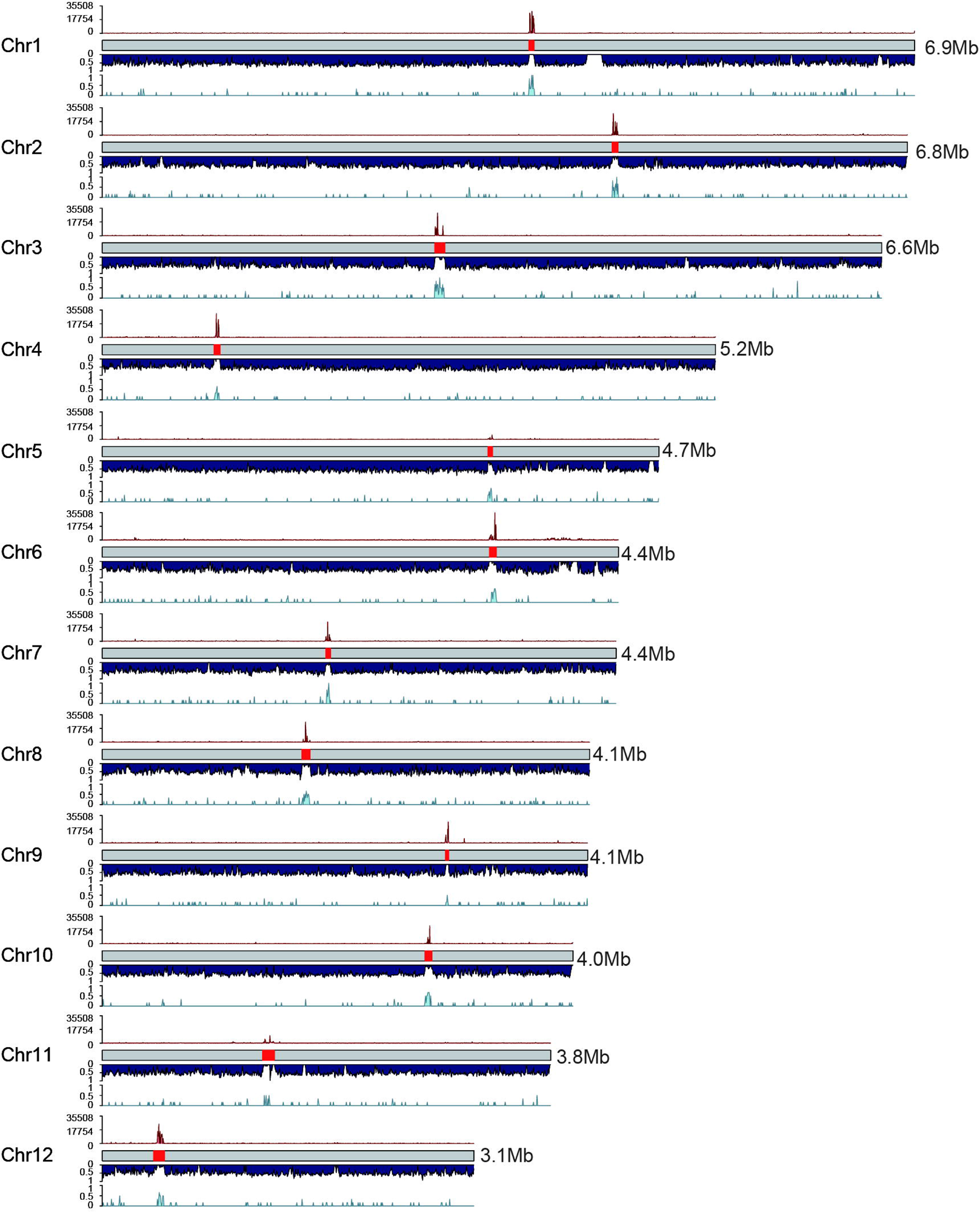
**CENH3 binding of the centromeres in *Chlorella sorokiniana* NS4-2**. Schematic overview of the chromosomes of *C. sorokiniana* NS4-2 showing, from top to bottom per chromosome, the normalized CENH3 ChIP-seq read coverage, predicted centromere regions, gene density, and LTR/Copia element density. Densities were calculated in 5 Kb windows, and normalisation conducted in 1 Kb bins with 3 Kb smoothing.

### Centromere methylation analysis in two *Chlorella* species

In higher plants, such as *Arabidopsis thaliana* and *Zea mays*, satellite repeats that are associated with chromatin containing CENH3 exhibit lower levels of methylation when compared to the non-CENH3 centromeric regions [10, 28]. These findings imply that DNA methylation is a significant factor in the epigenetic demarcation of centromeric chromatin. However, it is currently unclear whether centromeres in green algae exhibit similar methylation patterns. We utilized ccsmeth [29], a deep learning-based circular consensus sequencing (CCS) sequencing methylation calling software, to analyze the centromere methylation in the *Chlorella sorokiniana* NS4-2 genome (**Fig. 3**). Interestingly, we found that the centromeric region of chromosome 1 in *Chlorella sorokiniana* NS4-2 contains hypomethylated intervals (**Fig. 3A**). We more closely examined the regions 50 Kb up- and downstream of the centromere and observed a negative correlation between the hypomethylated regions and CENH3-ChIP signals (**Fig. 3A and Supplementary Fig. S12**). We then divided the genome into four categories – centromeric regions, non-centromeric regions, CENH3-ChIP peak regions in centromeric regions, and non-ChIP peak regions in centromeric regions – and conducted a statistical analysis of methylation frequencies. Our analysis revealed that centromeric regions exhibited significantly lower methylation levels compared to non-centromeric regions, which we attributed primarily to the low methylation status of the CENH3-ChIP peak regions (**Fig. 3B**). For *Chlorella pyrenoidosa* DBH, we employed HiFi and ONT sequencing to detect methylation status, and similarly identified regions of low methylation within the centromeres (**Supplementary Fig. S13**). We observed the same phenomenon of centromeric low methylation in *Chlorella sorokiniana* NS4-2 (**Fig. 3C**). These results indicate that despite the absence of tandemly repeated satellite sequences in the centromeres of the two *Chlorella* genomes, their pattern of methylation nevertheless resembles that of higher plants. This suggests that despite comparatively rapid centromeric evolution, their sequences retained a conserved epigenetic inheritance.

**Fig. 3.**
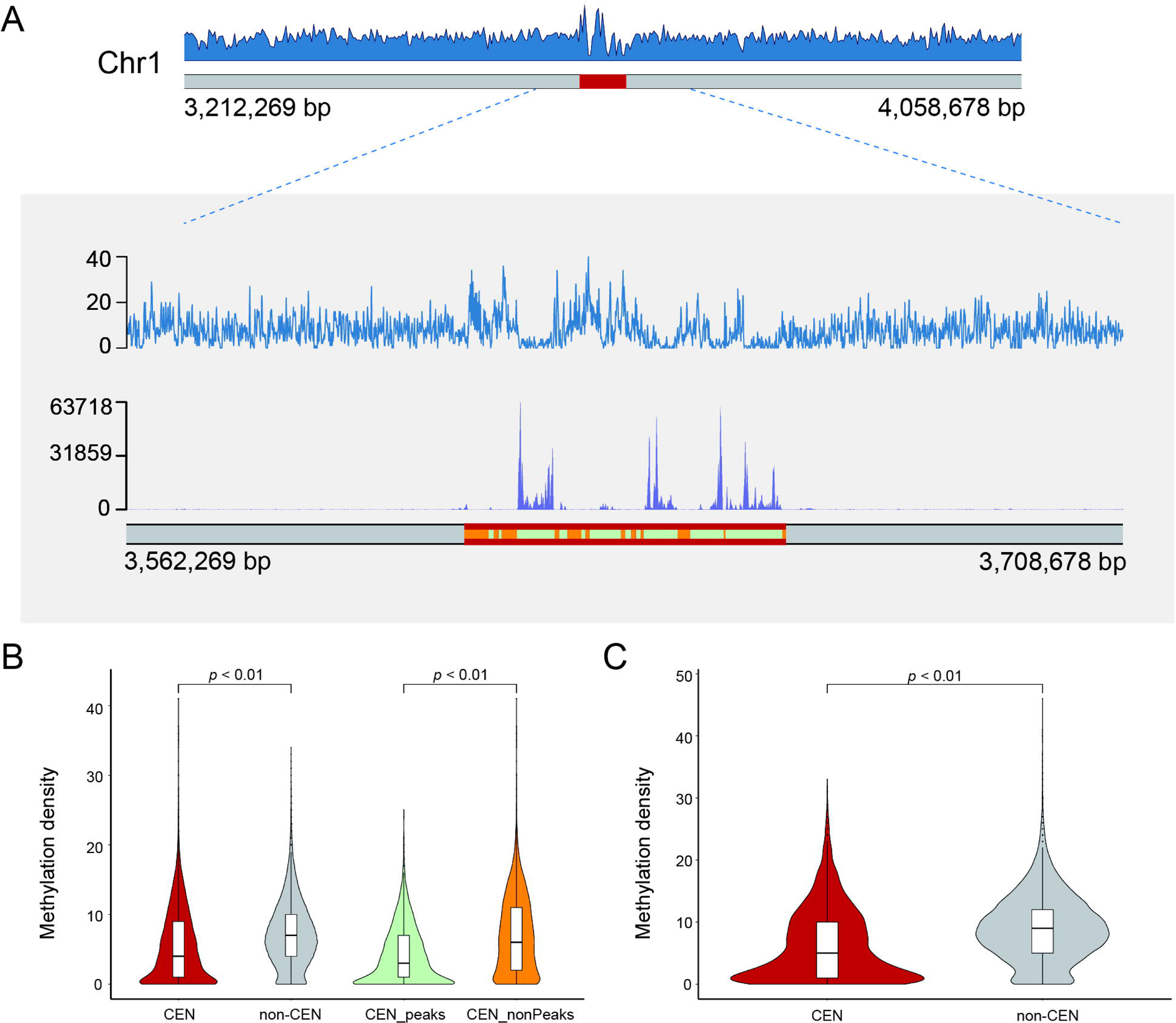
Centromeric methylation features of *Chlorella sorokiniana* NS4-2. and *Chlorella pyrenoidosa* DBH. (A) Methylation density distribution of *C. sorokiniana* NS4-2 chromosome 1. The red area represents the centromere. The close-up view shows, from top to bottom, the distribution of methylation density, CENH3 ChIP-seq signals, and the location of centromeric regions (parallel red band). Within the centromeric region, ChIP-seq peaks called by MACS2 are shown in light green and non-peaks in orange; **(B)** Box and violin plots showing methylation density in *C. sorokiniana* NS4-2 for centromeric (CEN) and non-centromeric (non-CEN) regions, and for peaks (CEN_peaks) and non-peaks (CEN_nonPeaks) within the former; **(C)** Box and violin plots showing methylation density in *C. pyrenoidosa* DBH for CEN and non-CEN regions. The statistical analysis was conducted via the Wilcoxon test.

### Centromere size and composition in green algae

We next selected six high-quality green algal assemblies for a compositional analysis of their putative centromeres: *Chlamydomonas reinhardtii* CC-4532 (v. 6.1) [24], *Chromochloris zofingiensis* [26], *Coccomyxa subellipsoidea* C-169 (v. 3.0) [25], *Chlorella* sp. SLA-04 [30], and the two *Chlorella* species assembled in this study (**Fig. 4A**). We used the red alga *Cyanidioschyzon merolae* (which has short low-GC centromeric regions) [31, 32] as an outgroup when constructing the phylogenetic tree of this lineage (**Fig. 4A**). For each of *Chlamydomonas reinhardtii*, *Chromochloris zofingiensis*, *Coccomyxa subellipsoidea* and *Chlorella* sp. SLA-04, we identified the potential centromeric loci using dot plots (**Supplementary Fig. S10 and Fig. 4C-H**) and prior knowledge [17, 25, 26], with average sizes of approx. 200 Kb, 27 Kb, 23 Kb, and 52 Kb, respectively (**Supplementary Table S4-S7**). We provided a detailed annotation of the centromere composition of each species in **Supplementary Tables S8 to S13**. In addition, we found a positive correlation between centromere and chromosome size for the six species, with a significant Pearson correlation coefficient of 0.87 (**Fig. 4B**). This fills a gap in the literature by providing data on green algae, which was previously lacking [33].

**Fig. 4.**
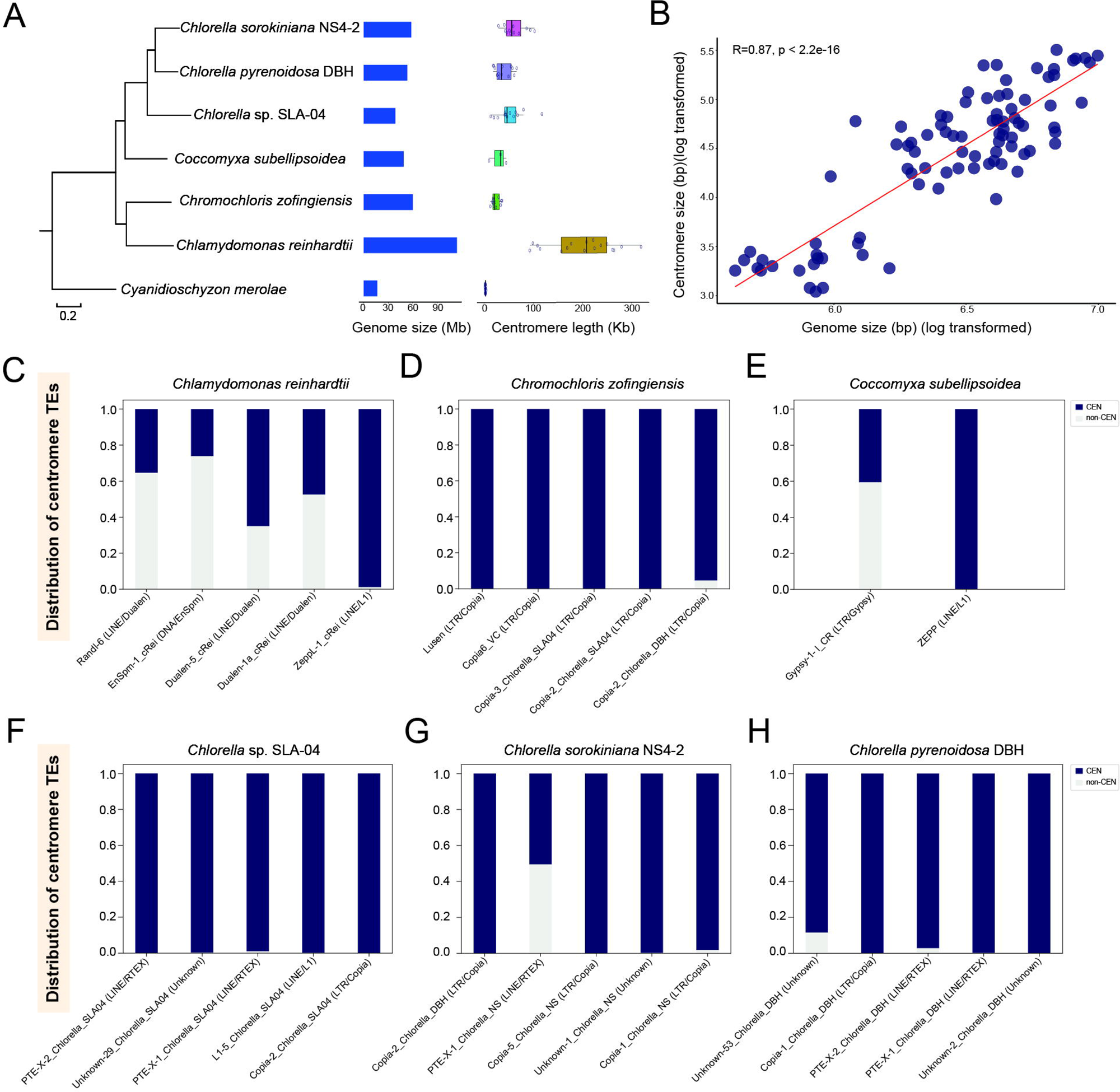
Centromeric size and composition in green algae. (A) Phylogenomic tree of six green algae species with the red algae *Cyanidioschyzon merolae* utilized as an outgroup. The figure on the right illustrates a comparison of genome size and centromere length among these selected species; **(B)** Relationship between the (log-transformed) lengths of chromosome and centromere; **(C-H)** The distribution of different repeat subfamilies in centromeres (CEN; dark blue) and across the genome (non-CEN; light gray) for *Chlamydomonas reinhardtii* **(C)**, *Chromochloris zofingiensis* **(D)***, Coccomyxa subellipsoidea* **(E)**, *Chlorella* sp. SLA-04 **(F)**, *Chlorella sorokiniana* NS4-2 **(G)** and *Chlorella pyrenoidosa* DBH **(H)**.

In the centromeres of both *Chlamydomonas reinhardtii* (**Fig. 4C and Supplementary Table S14**) and *Coccomyxa subellipsoidea* (**Fig. 4E and Supplementary Table S15**), we observed a substantial number of ZeppL-1_cRei (LINE/L1) and Zepp (LINE/L1) elements, respectively, consistent with previous reports [17]. Nearly all of these LINE/L1 elements are distributed in the centromere region, suggesting they are centromere-specific insertion (**Supplementary Table S8 and Supplementary Table S10**). Of note, however, is that although *Chromochloris zofingiensis* is closely related to *Chlamydomonas reinhardtii*, its centromeres primarily comprise members of the LTR/Copia, not LINE/L1, family (**Fig. 4D and Supplementary Table S16**).

The TE composition of the centromeres of each species are notably diverse. In *Chlorella* sp. SLA-04, the centromeres largely comprise Copia-2_Chlorella_SLA04 (LTR/Copia), L1-5_Chlorella_SLA04 (LINE/L1) and RTE-X-1_Chlorella_SLA04 (LINE/RTE-X) elements, at 37%, 22%, and 11% of the total, respectively (**Supplementary Table S11 and Supplementary Table S17**). Notably, the latter two elements are detected in all 13 centromeres of *Chlorella* sp. SLA-04 (**Supplementary Fig. S14**).

By contrast, in *Chlorella sorokiniana* NS4-2, the centromeres largely comprise Copia-1_Chlorella_NS (LTR/Copia) and Unknown-1_Chlorella_NS elements, at 50% and 21% of the total, respectively (**Supplementary Table S12 and Supplementary Table S18**), with both elements identified in all 12 centromeres (**Supplementary Fig. S15**). In *Chlorella pyrenoidosa* DBH, the largest centromeric component, at 25% of the total, is Unknown-2_Chlorella_DBH (**Supplementary Table S13 and Supplementary Table S19**), although RTE-X-2_Chlorella_DBH (LINE/RTE-X) elements, which represent another 20% of the total, were the only one found in all 13 centromeres (**Supplementary Fig. S16**). As centromeres are ‘cold spots’ for recombination [9, 34], it has been hypothesised that they could ‘protect’ TEs which insert there from being purged by purifying selection [35]. This would ultimately lead to the formation of TE-enriched regions specific to the centromeres, an interpretation consistent with our findings (**Fig. 4F-H**).

### Centromere evolution in green algae

Finally, we systematically correlated the primary constituents of each centromere (that is, their TEs) across the phylogenomic tree, an approach which allows us to examine the evolutionary trajectories of these sequences in different taxa (**Fig. 5A**). As LINE/L1 and LTR/Copia elements were found within both the Chlorophyceae and Trebouxiophyceae lineages (**Fig. 5A**), the implication is that these elements would also have resided within the centromeres of the most recent common ancestor of green algae.

**Fig. 5.**
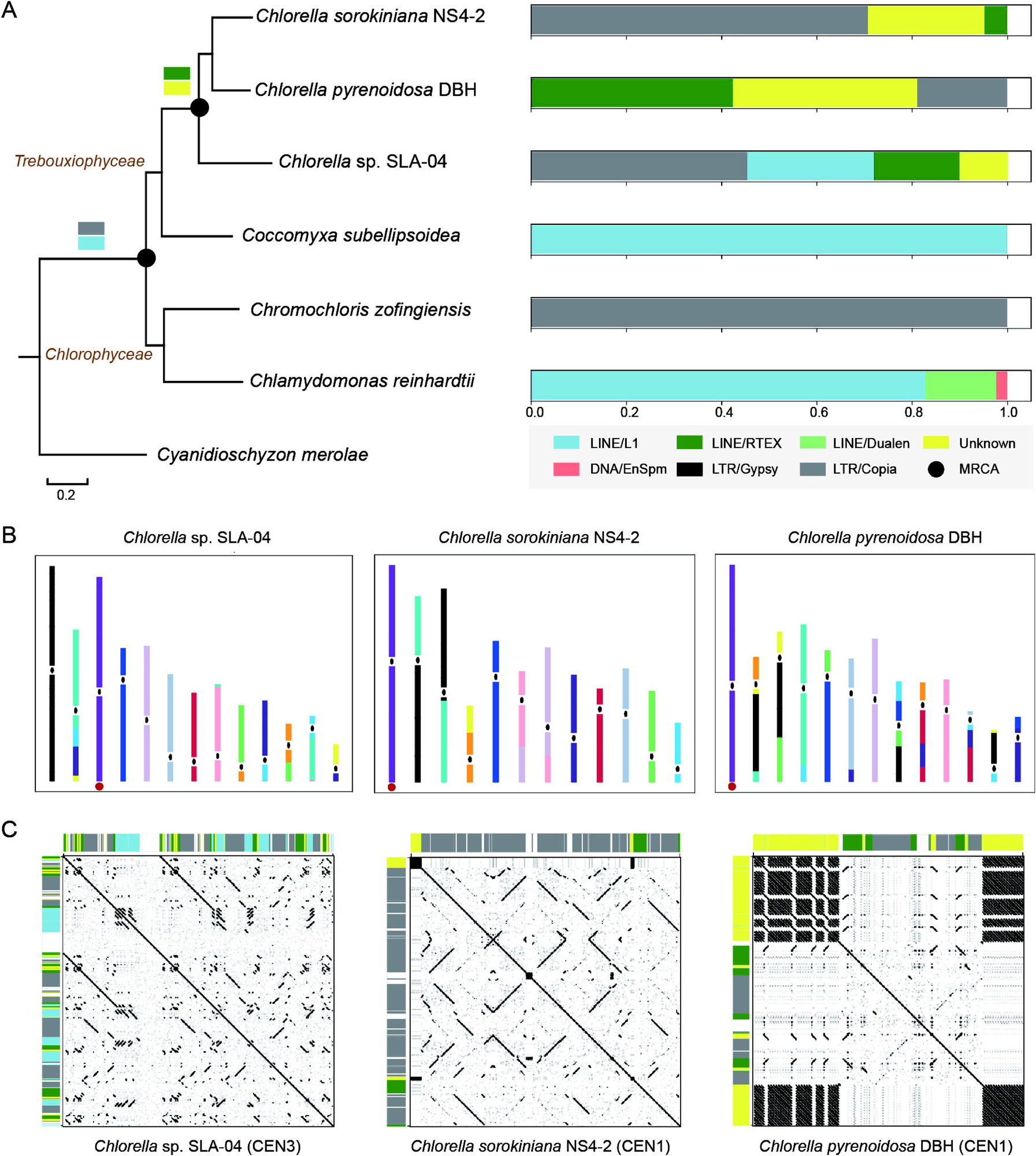
Evolution of centromeric transposable elements in green algae. (A) Predominant transposable elements (TEs) in the centromeres of green algal species. Different colors represent different TE families. MRCA represents the most recent common ancestor; **(B)** Synteny analyses of the 13 chromosomes of *Chlorella* sp. SLA-04, 12 chromosomes of *Chlorella sorokiniana* NS4-2 and 13 chromosomes of *Chlorella pyrenoidosa* DBH. The corresponding syntenic regions are marked using the same color. Approximate locations of putative centromeres are indicated by ellipses. The red circles represent the collinearity of the whole chromosomes; **(C)** Dot plot representation of the centromeres showing the collinearity of entire chromosomes from three *Chlorella* species, along with their TE composition. As with panel (A), different colors represent different TE families.

The *Chlorophyceae*-*Chlorella* divergence is estimated to have occurred approximately 827 MYA, while the *Coccomyxa*-*Chlorella* divergence is estimated to have occurred at around 544 MYA (http://www.timetree.org/). Our analyses of centromere composition suggest that for *Chlamydomonas reinhardtii*, *Chromochloris zofingiensis*, and *Coccomyxa subellipsoidea*, the centromeres comprise a primary TE component, seemingly a singular composite of many inserted elements, but that for *Chlorella* species, multiple diverse TE families are maintained within the centromeres. To explore this further, we performed collinearity analysis on the assemblies of three *Chlorella* species (*Chlorella* sp. SLA-04, *Chlorella sorokiniana* NS4-2, and *Chlorella pyrenoidosa* DBH), followed by illustrating the collinearity relationships and the putative centromeres (**Fig. 5B**). For example, chromosome 3 of *Chlorella* sp. SLA-04, chromosome 1 of *Chlorella sorokiniana* NS4-2, and chromosome 1 of *Chlorella pyrenoidosa* DBH exhibited strong collinearity (**Supplementary Fig. S17**), suggesting they share a common ancestor (**Fig. 5B**). We extracted the centromeres of these three chromosomes for dot plot analysis and found significant structural (that is, TE) differences among them (**Fig. 5C**). For instance, 88% of the centromere of *Chlorella sorokiniana* NS4-2 chromosome 1 comprised members of the LTR/Copia family, compared to 44% for *Chlorella* sp. SLA-04 chromosome 3 and 27% for *Chlorella pyrenoidosa* DBH chromosome 1. The implication of these results is that functional centromeres can vary greatly in their structure, and that the evolution of centromeric sequences is faster than that of non-centromeric ones.

## Discussion

Recent advances in long-read sequencing technologies have enabled the telomere-to-telomere (T2T) assembly of a number of genomes including the model plant *Arabidopsis thaliana* [10], the model green alga *Chlamydomonas reinhardtii* [23], and the model moss *Physcomitrium patens* [18]. To complement these, we employed HiFi reads, ONT ultra-long reads, and stringent assembly methods to generate T2T-quality assemblies for *Chlorella sorokiniana* NS4-2 and *Chlorella pyrenoidosa* DBH. These assemblies represent a significant improvement over previous genome assemblies, such as *Chlorella sorokiniana* UTEX1230 [36] and *Chlorella pyrenoidosa* FACHB-9 [37], and in that respect they may serve as valuable resources for future green algae research, in particular towards understanding the more complex regions of the genome, including the centromere.

In human centromeres, there are specific hypomethylated regions containing a higher concentration of the centromeric histone CENP-A, suggesting a functional role as kinetochore-binding sites [38, 39]. Interestingly, this study reveals that the same epigenetic phenomenon, which is conserved in higher plants [10, 28], is also present in green algae, suggesting a highly conserved epigenetic trait possibly among eukaryotes in general.

Centromeric satellite sequences typically have a monomer size of 100-200 base pairs, often 150 to 180 bp, a length sufficient to wrap a single nucleosome [4]. However, in plants, the origin of these monomers remains unclear. Analysis of ancestral crop species, such as *Gossypium raimondii* [40], an ancestor species of tetraploid cotton, and *Triticum monococcum* [16], the first domesticated wheat species, found that their centromeres were comprised largely of retrotransposons rather than long arrays of centromeric satellites, supporting the hypothesis that centromere repeats originated from non-random retrotransposon insertions [15]. We found that each of the green algae in this study had TE-rich centromeres, further supporting the hypothesis that the centromeric monomers of higher plants originated from TE insertions in their ancestors.

Nonetheless, there remain substantial variations in the composition of centromeric TEs among different taxa of green algae, likely reflecting differential TE insertion, expansion and purging. For example, *Chlamydomonas reinhardtii* and *Coccomyxa subellipsoidea* both harbor LINE/L1 elements within their centromeric regions although from different subfamilies [17]. Furthermore, the two *Chlorella* genomes sequenced in this study, despite belonging to the same genus, differ greatly in the composition and structure of their centromeres. These observations lend further support to the view that the centromere is the fastest-evolving part of the chromosome [41, 42].

In summary, we assembled two *Chlorella* species (*Chlorella sorokiniana* NS4-2 and *Chlorella pyrenoidosa* DBH) to T2T level, enabling a comprehensive analysis of their centromeric sequence composition and evolution. These findings contribute to our understanding of centromere evolution in green algae and, by extension, to their descendants.

## Materials and methods

### Culture conditions

*Chlorella pyrenoidosa* strain DBH (obtained from the Freshwater Algae Culture Collection at the Institute of Hydrobiology, Chinese Academy of Sciences) and *Chlorella sorokiniana* strain NS4-2 (isolated and purified from a water sample taken from Xueshi Lake at Shanghai Normal University) were grown mixotrophically in a tris-acetate-phosphate (TAP) medium. Both species were cultivated in Erlenmeyer flasks under continuous growth light conditions (30-40 μmol photons m^−2^ s^−1^) at a temperature of 25 °C, with agitation at 100 rpm, for maintenance purposes.

### Karyotyping of the two *Chlorella* species

Culture samples were harvested by centrifugation at 12,000 rpm for 2 min and then washed sequentially with sterilized BG-11 medium and distilled water. The resulting algal pellets were suspended in a 2 mM 8-hydroxyquinoline solution and kept in darkness at 4 °C for 4 h. Subsequently, the algae pellets were fixed with a solution of 3 parts ethanol to 1 part acetic acid (v/v) for 24 h to ensure fixation of the algae cells in the metaphase of mitosis. After fixation, the cells were washed three times with distilled water and then subjected to enzymatic digestion. The enzymatic solution contained 2.5% (w/w) cellulose Onozuka R 10 (Yakult Pharmaceutical, Tokyo, Japan) and 2.5% (w/w) pectinase Y23 (Yakult Pharmaceutical, Tokyo, Japan), and the digestion process lasted for 18 h at 37 °C. Following digestion, the cells were homogenized at 15,000 rpm for 2 min and then centrifuged. The resulting pellets were resuspended in 100 µL of acetic acid and vortexed for 20 sec. The cell suspension was dropped onto a sterile slide and allowed to air dry. Finally, the cells were stained with an improved carbol fuchsin solution and examined under a microscope to capture images of the metaphase stage.

### Genomic DNA extraction

Samples were collected, and high molecular weight genomic DNA was prepared by the CTAB method followed by purification with the QIAGEN® Genomic kit (Cat#13343, QIAGEN) for regular sequencing, according to the standard operating procedure provided by the manufacturer. DNA degradation and contamination of the extracted DNA was monitored on 1% agarose gels. DNA purity was then assessed using a NanoDrop One UV-VIS spectrophotometer (Thermo Fisher Scientific, USA), of which OD 260/280 ranged from 1.8 to 2.0 and OD 260/230 from 2.0 to 2.2. DNA concentration was subsequently measured using a Qubit 4.0 fluorometer (Invitrogen, USA).

### HiFi sequencing and genome assembly

SMRTbell libraries were prepared following the standard protocol of PacBio, using 15 Kb preparation solutions from Pacific Biosciences, USA. The library preparation steps were conducted according to the method described in [10]. Sequencing was performed on a PacBio Sequel II instrument with Sequencing Primer V2 and Sequel II Binding Kit 2.0 at Xi’an Haorui Genomics Technology Co., Ltd. (Xi’an, China). For *Chlorella sorokiniana* strain NS4-2, a total of 25.87 Gb of HiFi reads were generated, providing a coverage of 438×. The N50 of the reads was 15,694 bp. Regarding the *Chlorella pyrenoidosa* strain DBH, a total of 23.74 Gb of HiFi reads were generated, resulting in a coverage of 308×. The N50 of the reads was 13,338 bp. The HiFi reads were assembled using hifiasm v. 0.16.1-r375 [6] with default parameters. The gfatools (https://github.com/lh3/gfatools) was used to convert GFA-format sequence graphs to FASTA format.

### ONT sequencing and genome assembly

For the ultra-long Oxford Nanopore sequencing library, approximately 8‒10 µg of genomic DNA was selected (>50 kb) with the SageHLS HMW library system (Sage Science, Beverly, MA), and then processed using the ligation sequencing 1D kit (SQK-LSK109, Oxford Nanopore Technologies, Oxford, UK) according to the manufacturer’s instructions. DNA libraries (approximately 800 ng) were constructed and sequenced using a PromethION (Oxford Nanopore Technologies, Oxford, UK) at the Genome Center of Grandomics (Wuhan, China). A total of 22.12 Gb ONT long reads with 368× coverage (read N50: 59,282 bp) were generated. To assemble these into contigs, the long-read assembler NextDenovo v. 2.0 [21] was used with parameters ‘read_cutoff = 5k’ and ‘seed_cutoff = 132,382’. Contigs were then polished using Nextpolish v. 1.3.1 [43] with parameters ‘hifi_options = -min_read_len 10k -max_read_len 45k’ and ‘hifi_minimap2_options = -x asm20’.

### Hi-C sequencing and scaffolding

A Hi-C library was generated from cross-linked chromatins of algal cells using a standard Hi-C protocol, and then sequenced using the NovaSeq 6000 platform. The Hi-C sequencing data, as outlined in Table S1, were utilized to construct a Hi-C contact map. The construction of the contact map was performed using Juicer v. 1.5 [44] with parameter ‘-s DpnII’. The resulting contact map was manually inspected and verified using Juicebox v. 1.11.08 [45].

### Evaluation of the putative centromeres

To assess the assembly structure of the predicted centromeric regions, we employed VerityMap [27]. First, the PacBio HiFi reads were aligned to the assembly using minimap2 v. 2.20-r1061 [46] with the ‘-x map-hifi’ parameter. Reads aligning to the centromeric regions were extracted using SAMtools v. 1.9 [47]. These extracted HiFi reads were then input into VerityMap using the following parameters: ‘--careful -- reads HiFi reads -d hifi’.

### Genome comparisons

The synteny relationships among the chromosomes were estimated using MCScanX [48] with parameter ‘-s 10’ and visualized using Circos v. 0.69-8 [49]. Genomic alignment dot plot of chromosomes was generated using Gepard v. 2.1 [50]. To assess genome completeness, we used BUSCO v. 3.0.2 with the dataset ‘chlorophyta_odb10 v. 4’ [51]. Chromosome painting among three species of the genus *Chlorella* was performed using IAGS [52] with the GMP model.

### Genome annotation

For the *Chlorella sorokiniana* NS4-2 and *Chlorella pyrenoidosa* DBH genome annotations, we initially predicted the *de novo* gene structure using GeneMark-ES with the parameters ‘--ES’ [53]. Subsequently, we employed the ‘chlorella gene model’ to train Augustus v. 3.3 [54]. To identify protein-coding genes, we utilized the MAKER2 pipeline [55], which integrated the gene models predicted from GeneMark-ES and Augustus, as well as the protein sequences from four closely related organisms: *Chlamydomonas reinhardtii* (GenBank accession: GCF_000002595.2) [56], *Chlorella variabilis* (GCF_000147415.1) [57], *Chlorella sorokiniana* strain 1602 (GCA_002245835.2) [58], and *Micractinium conductrix* (GCA_002245815.2) [58]. For the identification of repeat sequences, we employed RepeatMasker v. 4.0.7 (http://www.repeatmasker.org) and RepeatModeler v. 2.0.1 [59]. The TE-algae library used to annotate the selected species consists of three parts: the *de novo* library of three *Chlorella* species from RepeatModeler, a library from a previous study [17], and the *Chlorophyta* Repbase library (https://www.repeatmasker.org/) (**Supplementary Dataset S1**). The TE-algae library underwent redundancy reduction using cd-hit [60]. The karyoplots shown in this study were generated using the KaryoploteR package in R [61].

### Phylogenetic tree construction

The phylogenetic trees depicted in Fig. 4 and Fig. 5 were generated utilizing OrthoFinder v. 2.2.7 with the default parameters [62]. Supplementary Fig. S1’s phylogenetic tree was constructed using MEGA v. 7 software and employed the Kimura 2-parameter model, accompanied by 1000 bootstrap replicates [63].

### PacBio DNA methylation analysis

Methylation for *Chlorella sorokiniana* NS4-2 and *Chlorella pyrenoidosa* DBH was inferred with ccsmeth v.0.4.1 [29], using kinetic features from PacBio CCS reads. Methylation of CCS reads was predicted using ‘ccsmeth call_mods’ with the model ‘model_ccsmeth_5mCpG_call_mods_attbigru2s_b21.v2.ckpt’. Then, we used ‘ccsmeth align_hifi’ to align the reads to their respective genome. The methylation frequencies were calculated by using ‘ccsmeth call_freqb’ with the aggregate model ‘model_ccsmeth_5mCpG_aggregate_attbigru_b11.v2p.ckpt’.

### ONT DNA methylation analysis

Nanopolish v. 0.13.2 with the parameters ‘call-methylation --methylation cpg’ was used to measure the frequency of CpG methylation. The ONT reads were aligned to whole-genome assemblies using minimap2 [44] with default parameters. The script ‘calculate_methylation_frequency.py’, provided in the methplotlib package [64], was then used to generate the methylation frequency. Methylation density is defined as the number of occurrences with a methylation frequency greater than 0.8 within genomic sliding windows (100-base pair window advanced at 50-base pair increments).

### CENH3 ChIP-seq

We first used BLASTP to identify the amino acid sequence of CENH3 in *Chlorella sorokiniana* NS4-2, with the reference sequence HTR12 from *Arabidopsis thaliana* (Genbank accession: NP_001322867.1), and the ‘-evalue’ parameter set to 1e-30. We selected the coding gene NS|00958 as the unique matching result for the CENH3 sequence of the *Chlorella sorokiniana* NS4-2 species for subsequent antibody preparation. An antigen with the peptide sequence ‘MARTKQTKPSPSKRGA’ corresponding to the N terminus of NS4-2_CENH3 was used to produce the antibody (Beijing QiWei YiCheng Tech Co., Ltd.).

Shanghai Jiayin Biotechnology Co., Ltd conducted ChIP assays following a modified version of the standard crosslinking ChIP protocol. In summary, the culture sample was harvested by centrifugation at 12,000 rpm for 2 min and sequentially washed with sterilized BG-11 and distilled water. Subsequently, approximately 4g of algae pellet was obtained and ground in liquid nitrogen. The ground sample was fixed with 1% formaldehyde at room temperature for 15 min, followed by a 5-minute treatment with 0.125 M glycine. After fixation, the sample was washed, resuspended in a lysis buffer, and sonicated to generate DNA fragments. Following sonication, immunoprecipitation was carried out using the custom antibody NS4-2_CENH3. The immunoprecipitated complex was washed, and DNA was extracted. DNA purification was performed using the Universal DNA Purification Kit (#DP214). A ChIP-seq library was prepared by employing the ChIP-Seq DNA sample preparation kit (NEBNext® UltraTMII DNA), adhering to the manufacturer’s instructions. The extracted DNA was ligated to specific adaptors and subsequently subjected to deep sequencing on the Illumina Novaseq 6000 platform in 150bp paired-end mode. The raw reads were trimmed using the Trimmomatic program [65] with default parameters. The ChIP/input paired-end clean reads were mapped to the *Chlorella sorokiniana* NS4-2 genome using BWA-mem [66] with default parameters. We determined the ChIP/input coverage ratio by employing the DeepTools function ‘bamCompare’, with a threshold of MAPQ ≥ 30. The output format was in BigWig. We carried out peak calling using MACS2 [67] with the parameters ‘-t ChIP.bam -c input.bam -f BAM -- outdir chipseq -n chipseq -B --nomodel --keep-dup 1 -g 58000000’.

### Ethical approval and consent to participate

Not applicable.

### Consent for publication

Not applicable.

### Availability of data and materials

The whole genome sequence data reported in this paper have been deposited in the Genome Warehouse in National Genomics Data Center, Beijing Institute of Genomics, Chinese Academy of Sciences / China National Center for Bioinformation (GWH: GWHBKBA00000000 for *Chlorella sorokiniana* NS4-2; GWH: GWHBKBB00000000 for *Chlorella pyrenoidosa* DBH) and is publicly accessible at https://ngdc.cncb.ac.cn/gwh. The genome annotations of *Chlorella sorokiniana* NS4-2 and *Chlorella pyrenoidosa* DBH have been deposited in Figshare (10.6084/m9.figshare.24503881). The raw data for the PacBio HiFi reads, ONT long-reads, Hi-C Illumina short reads and ChIP-seq data have been deposited in the Genome Sequence Archive at the National Genomics Data Center, Beijing Institute of Genomics, Chinese Academy of Sciences / China National Center for Bioinformation (GSA: CRA007945), and are publicly accessible at https://ngdc.cncb.ac.cn/gsa.

## Authors’ contributions

K.Y. and W.M. conceived and designed the project. K.Y. and B.W. designed the experiments. B.W., Y.J., S.G., H.G. and X.Y. analyzed the results. B.W., Y.J., N.D., J.Y., S.B, S.G., W.H., S.W., and H.G. performed all the experiments and analyzed the data. B.W., Y.J. and N.D. prepared the figures. B.W., Y.J. and S.B. wrote the manuscript text. All authors reviewed and approved the manuscript.

