## Supplementary Figures 1-17 for "Near telomere-to-telomere genome assemblies of two *Chlorella* species unveil the composition and evolution of centromeres in green algae"

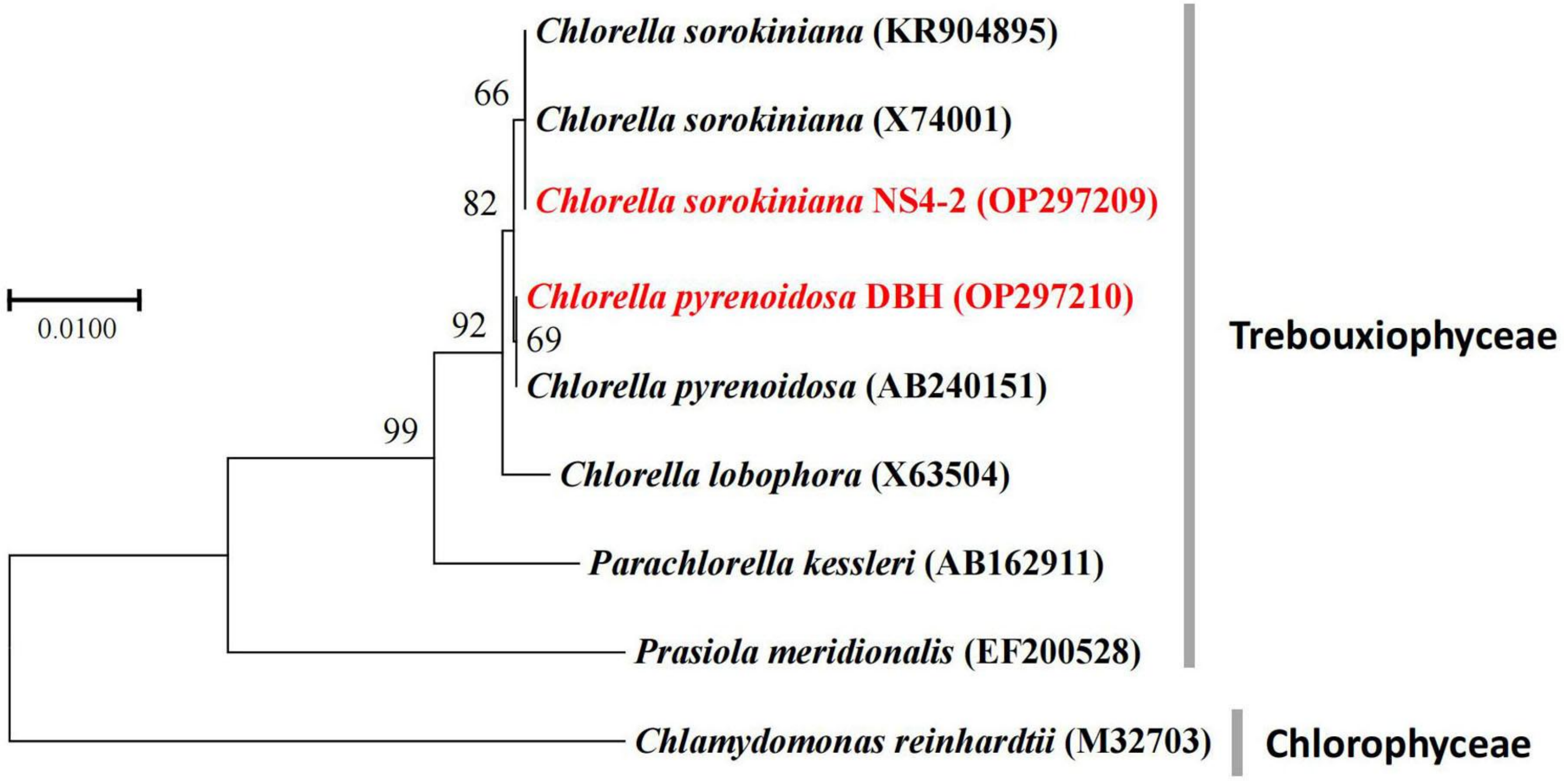


**Supplementary Fig. S1.** Maximum likelihood phylogenetic tree based on the analysis of 18S gene sequences. GenBank accession numbers in brackets.

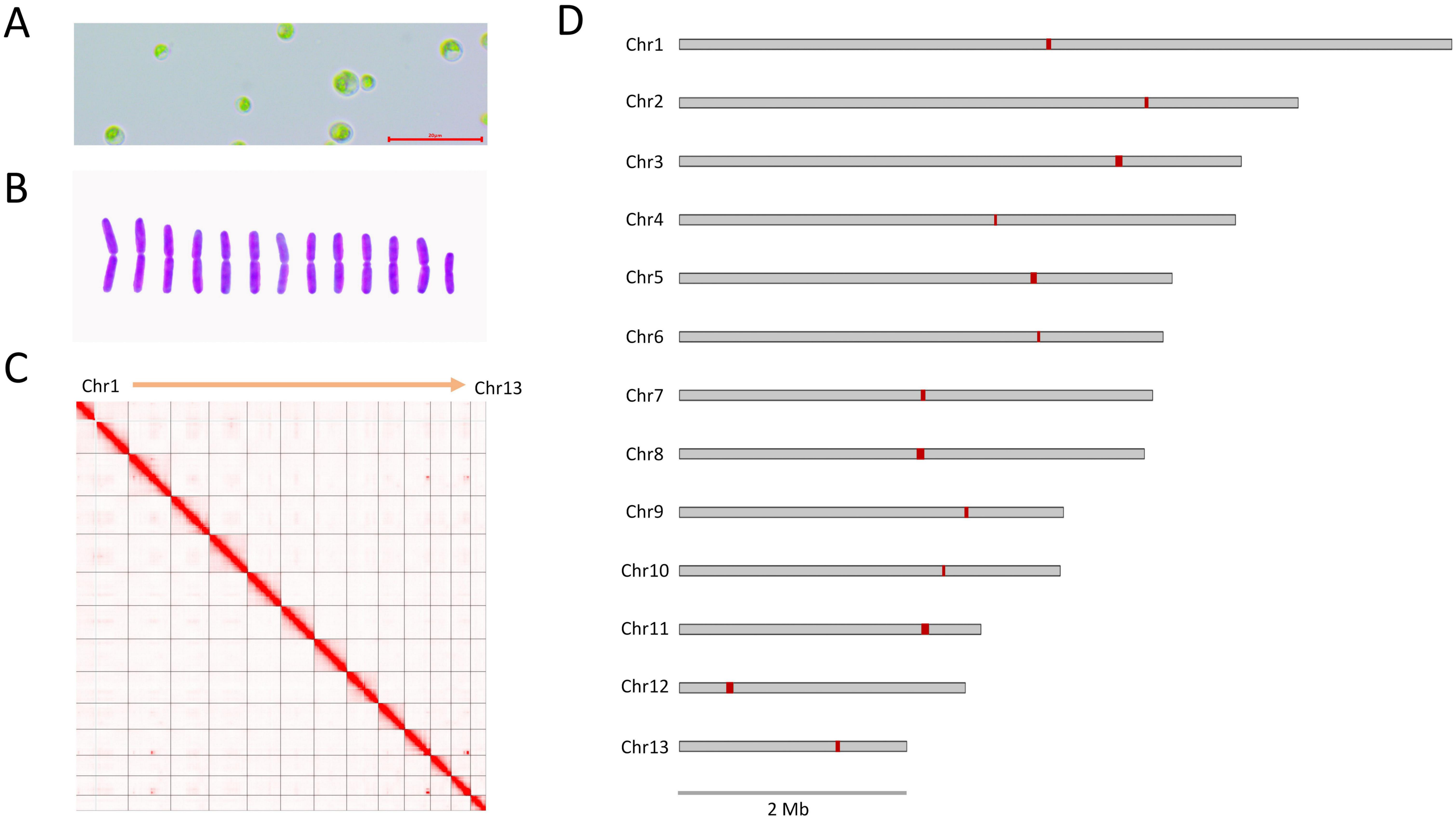


**Supplementary Fig. S2.** Near telomere-to-telomere genome assemblies and centromere identification of *Chlorella pyrenoidosa* strain DBH. **(A)** Micrograph of *C. pyrenoidosa* DBH, bar = 20 μm; **(B)** The karyotype diagram indicates that *C. pyrenoidosa* DBH has 13 chromosomes; **(C)** Hi-C contact map of the *C. pyrenoidosa* DBH T2T genome assembly (chromosome 1-13); **(D)** Schematic overview of the chromosomes of *C. pyrenoidosa* DBH showing the predicted centromere regions (red bands).

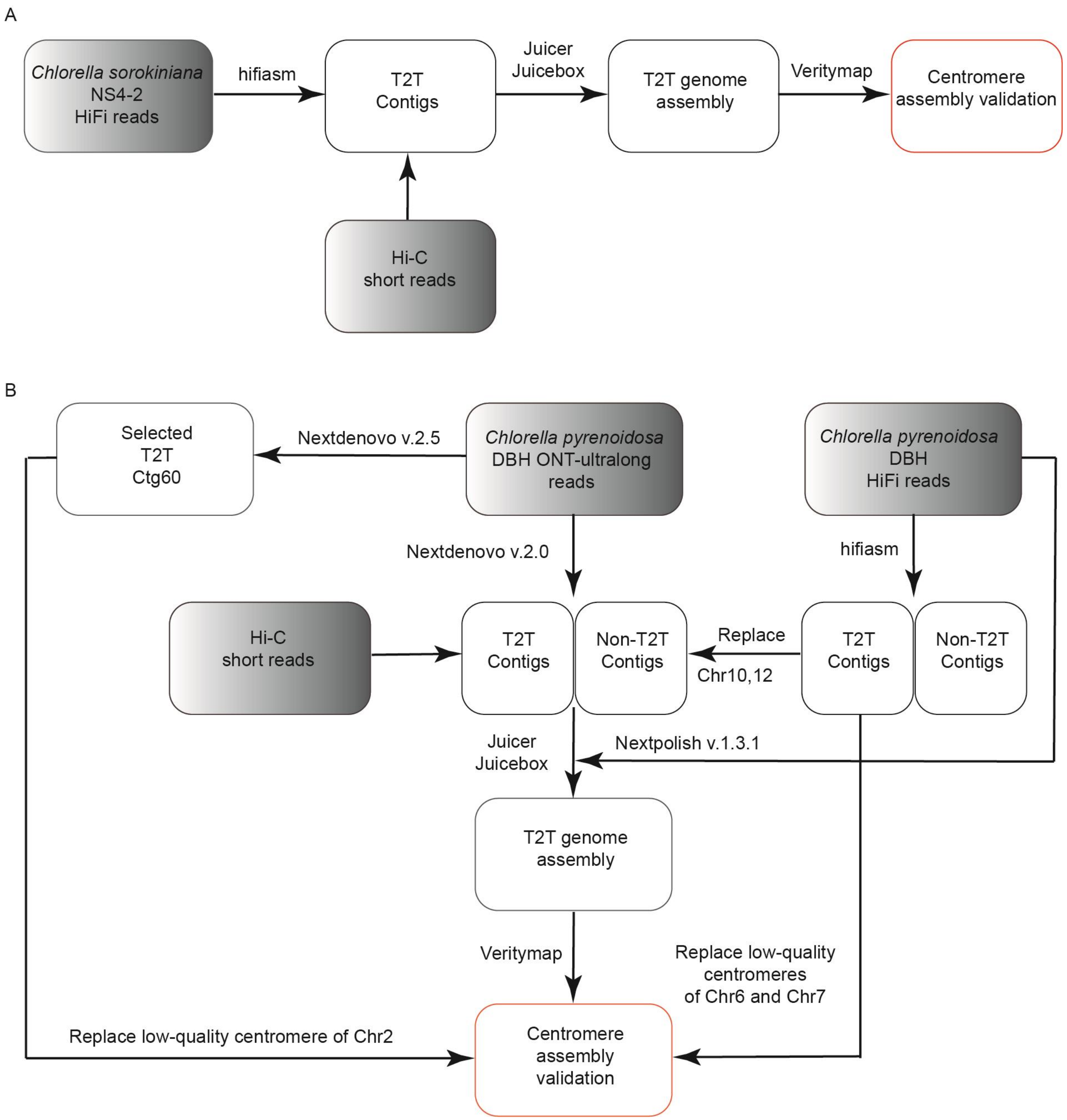


**Supplementary Fig. S3.** Overview of the telomere-to-telomere genome assembly pipelines. **(A)** *Chlorella sorokiniana* NS4-2 genome assembly; **(B)** *Chlorella pyrenoidosa* DBH genome assembly.

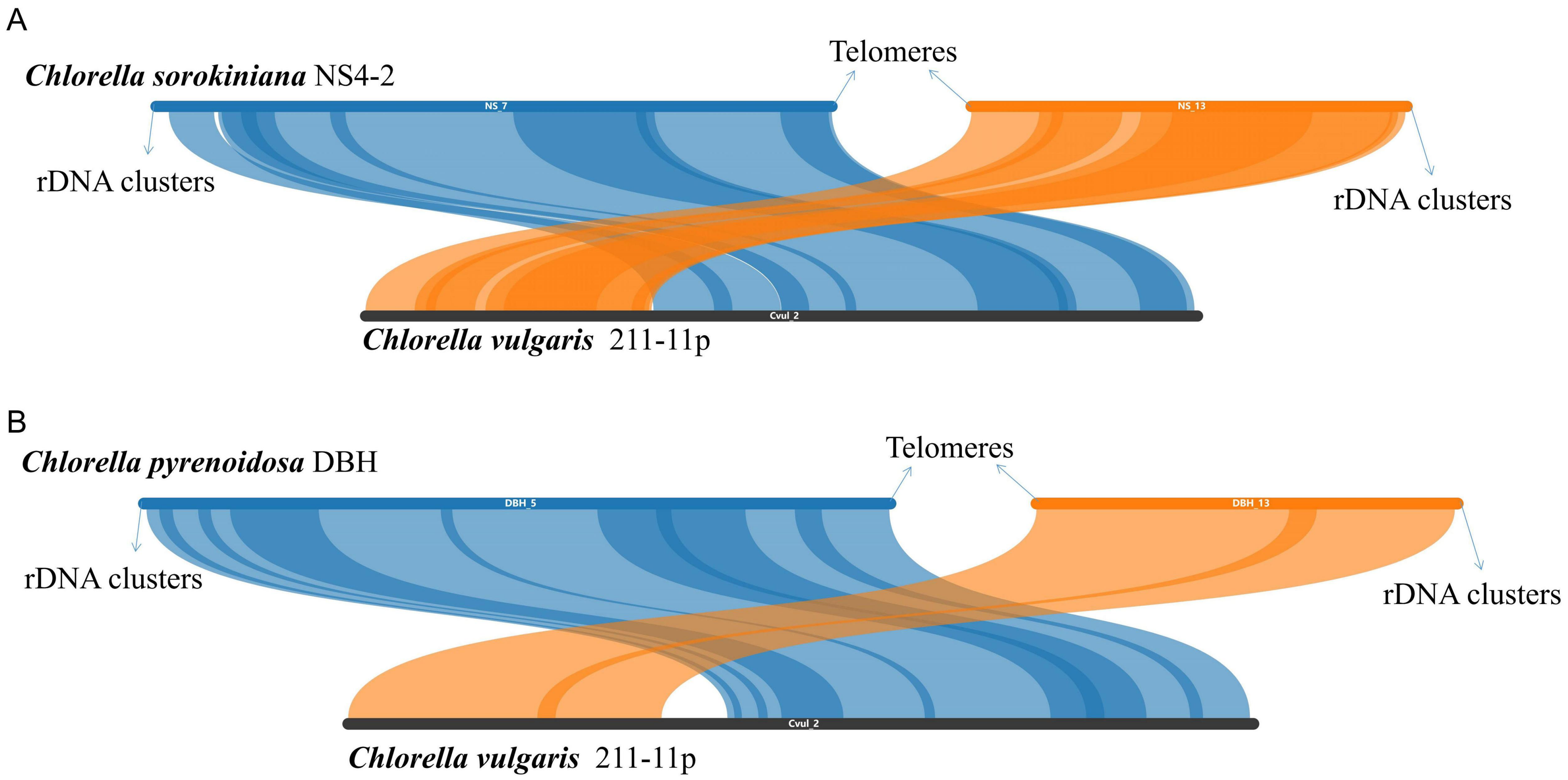


**Supplementary Fig. S4.** Illustration of collinearity between chromosomes. **(A)** Collinearity between two contigs of *Chlorella sorokiniana* NS4-2 and chromosome 2 of *Chlorella vulgaris*; **(B)** Collinearity between two contigs of *Chlorella pyrenoidosa* DBH and chromosome 2 of *Chlorella vulgaris*.

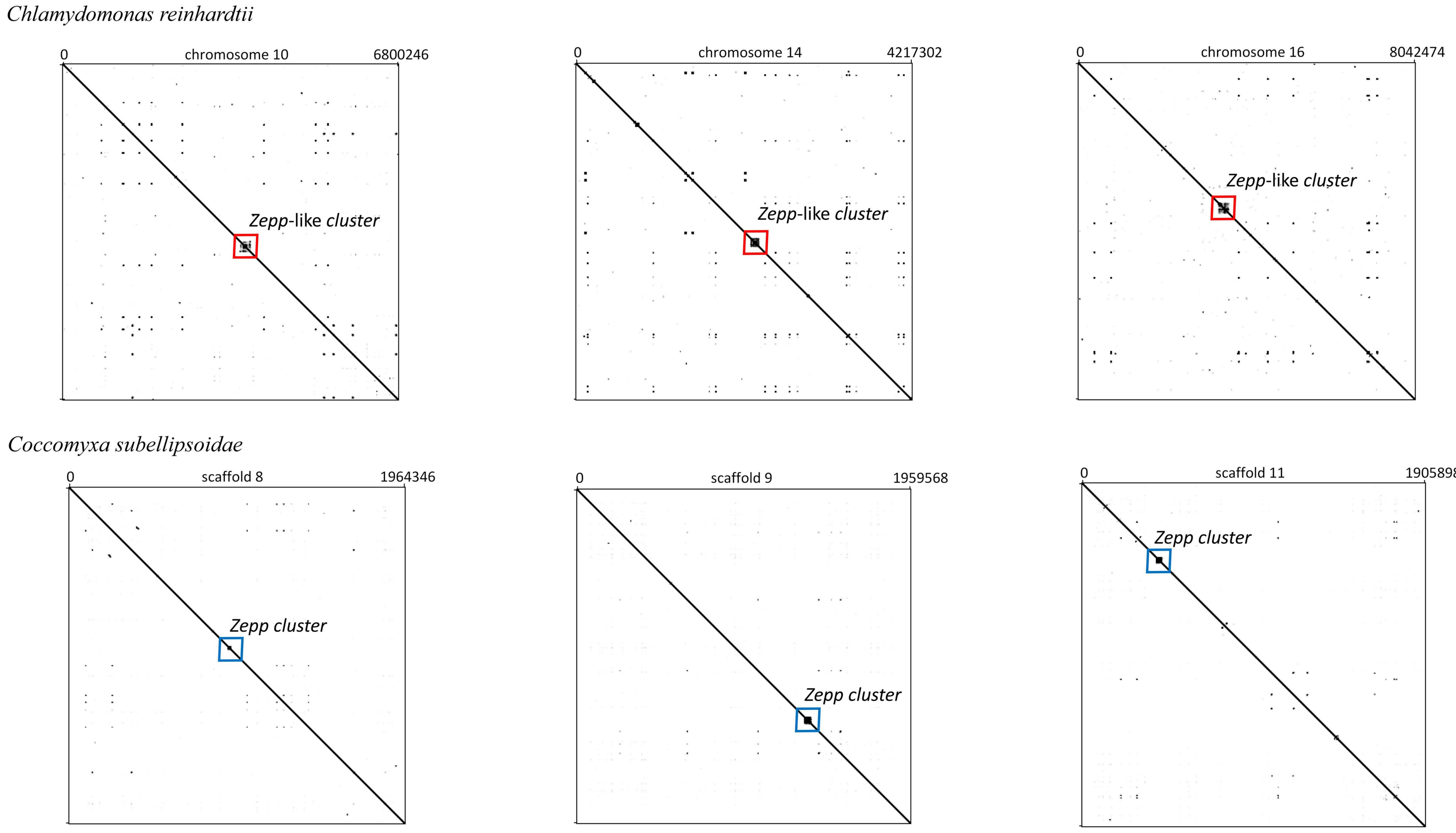


**Supplementary Fig. S5.** Self-dot plot showing potential centromeric features and regions of two previous assemblies, *Chlamydomonas reinhardtii* and *Coccomyxa subellipsoidae*.

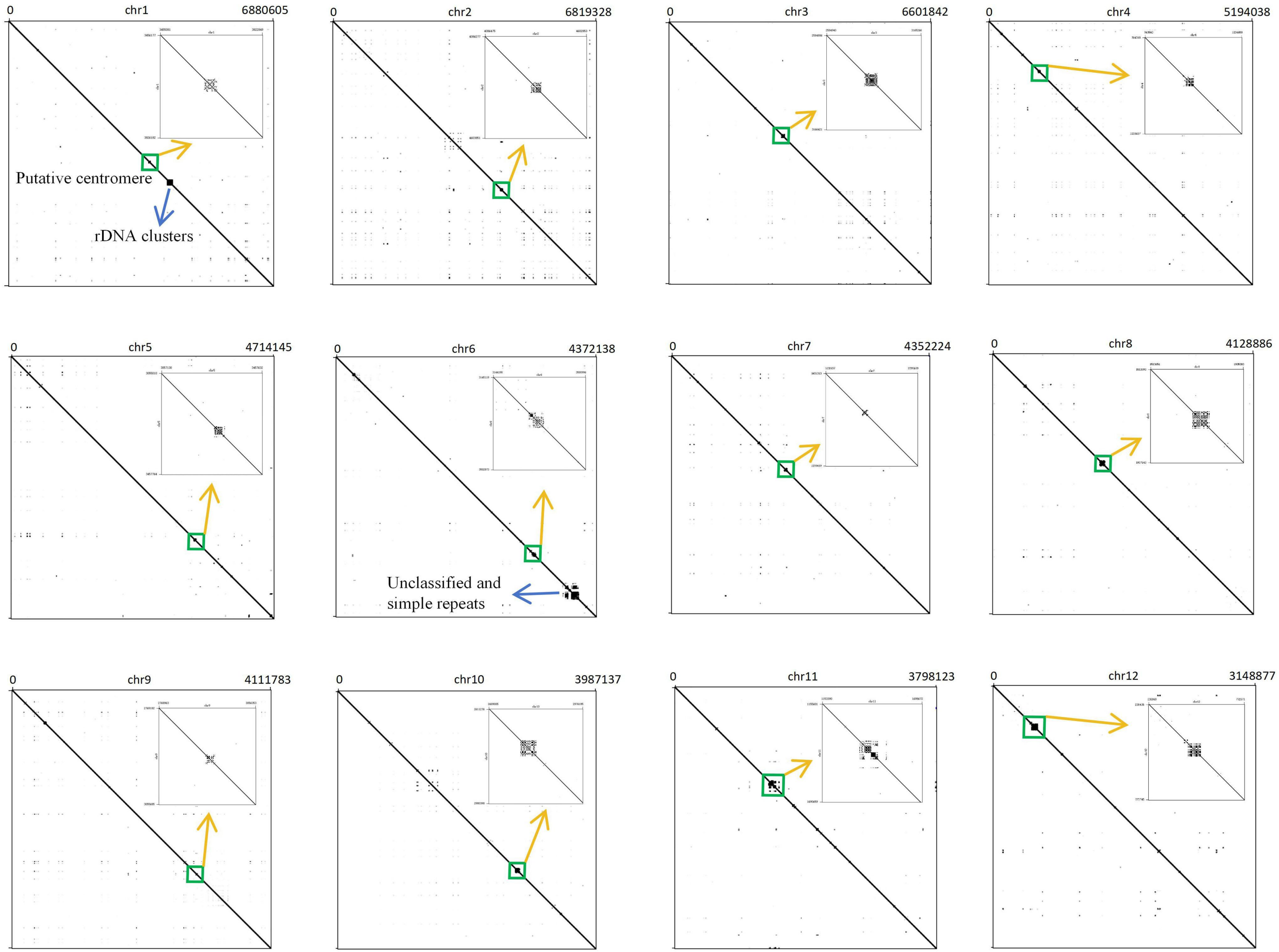


**Supplementary Fig. S6.** Self-dot plot of the 12 chromosomes of *Chlorella sorokiniana* NS4-2, with green boxes indicating potential centromeric regions.

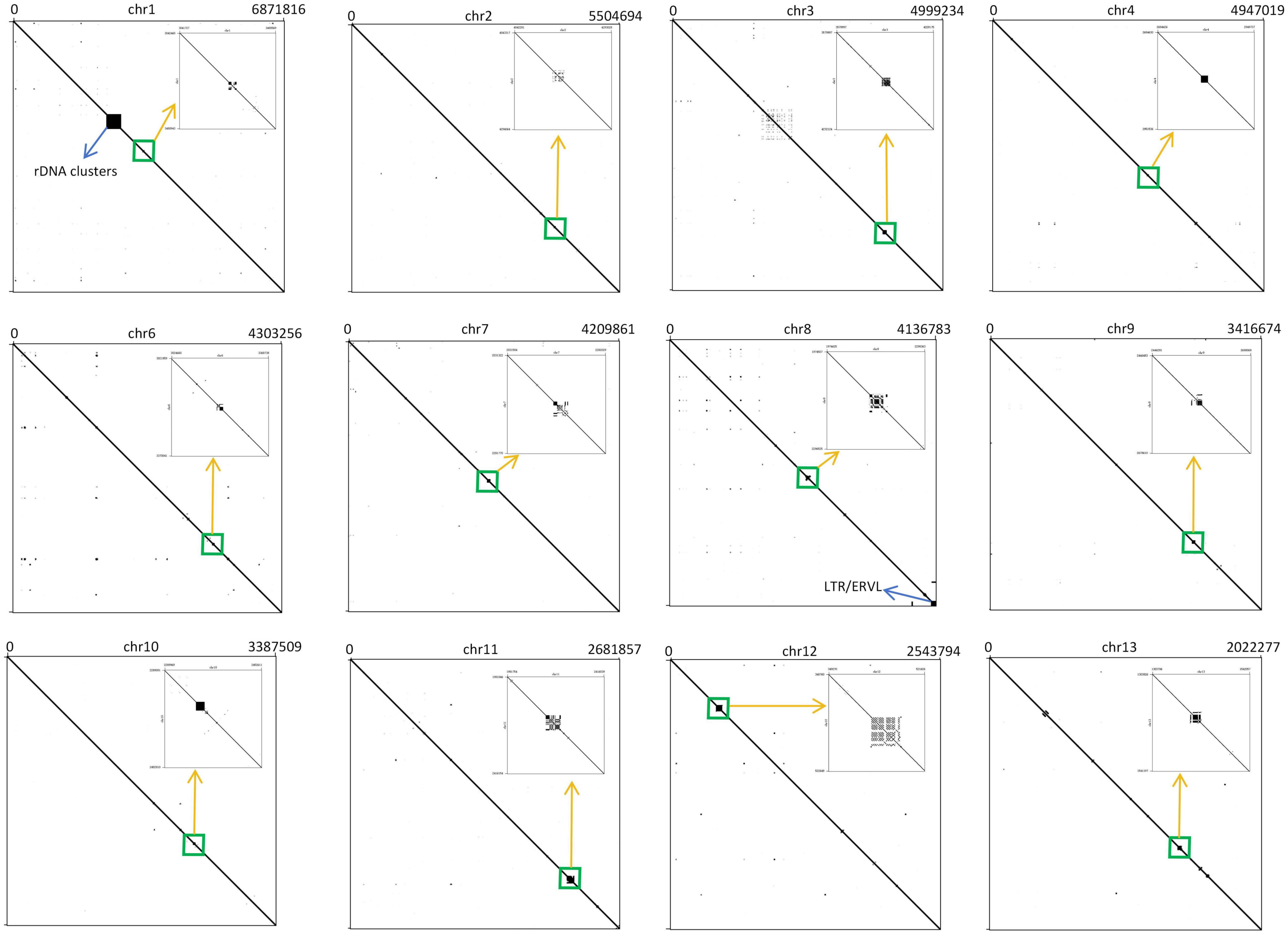


**Supplementary Fig. S7.** Self-dot plot of the 13 chromosomes of *Chlorella pyrenoidosa* DBH, with green boxes indicating potential centromeric regions.

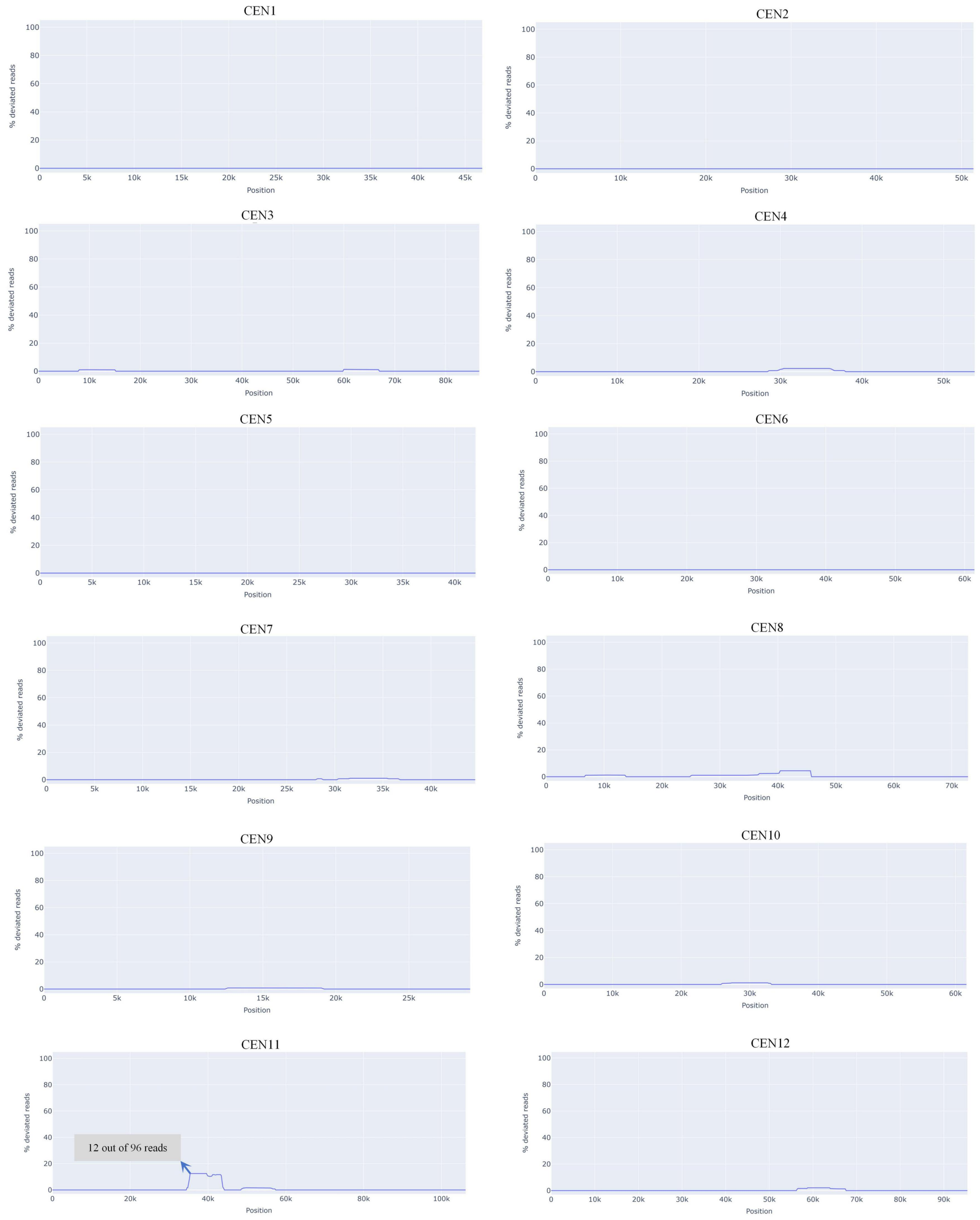


**Supplementary Fig. S8.** Evaluation of the 12 putative centromeric assemblies of *Chlorella sorokiniana* NS4-2 using VerityMap. The x-axis represents the position of the centromere, while the y-axis indicates the percentage of deviated reads, limited or negligible in each case. At most, only a low percentage (12.5%) of deviated reads were detected in the centromere of chromosome 11 (CEN11), which we interpret to mean there were no potentially large structural errors in the assembly.

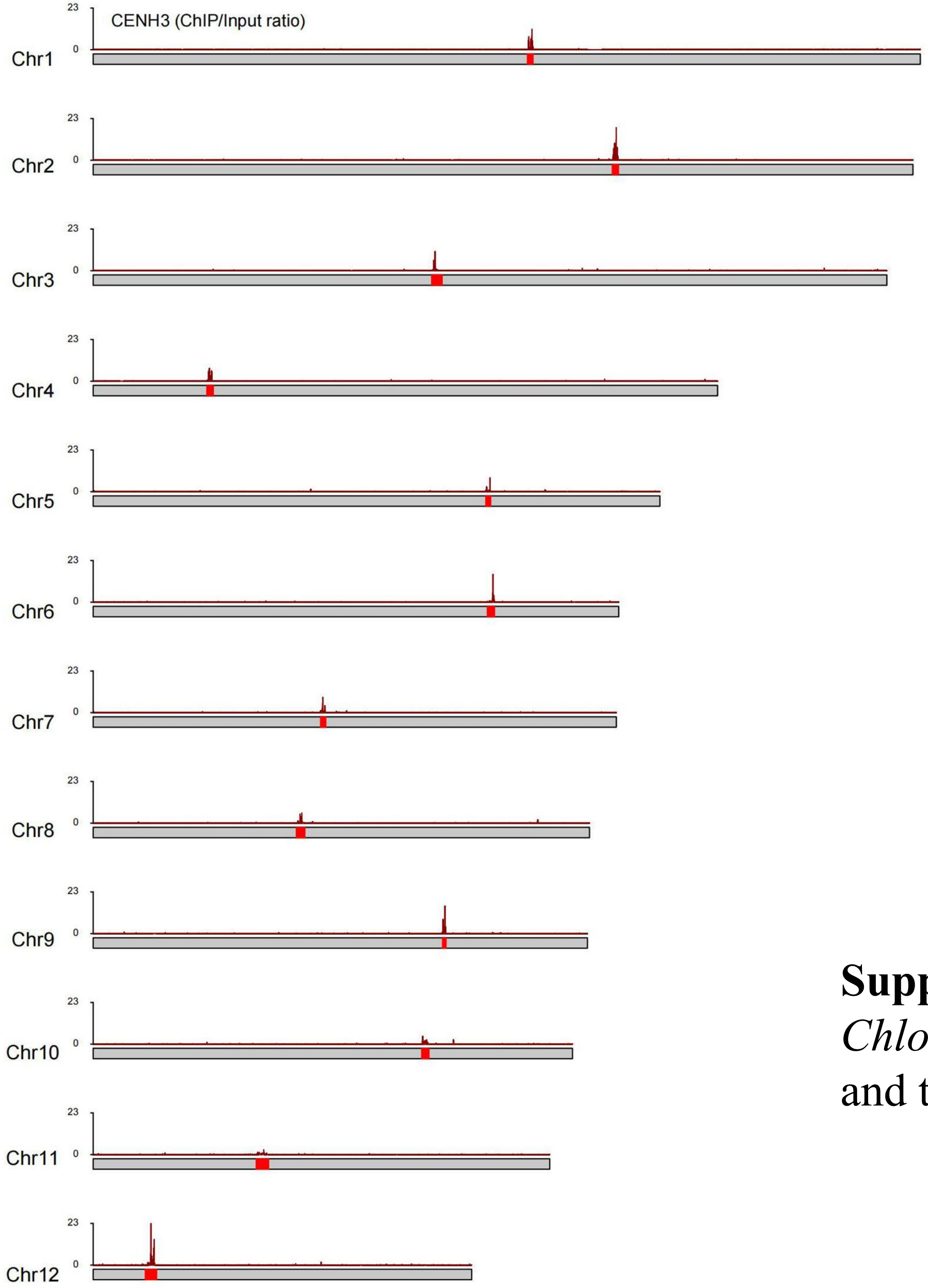


**Supplementary Fig. S9.** ChIP/Input ratio of the 12 chromosomes of *Chlorella sorokiniana* NS4-2. Schematic overview showing the ratio and the putative centromere regions (red bands).

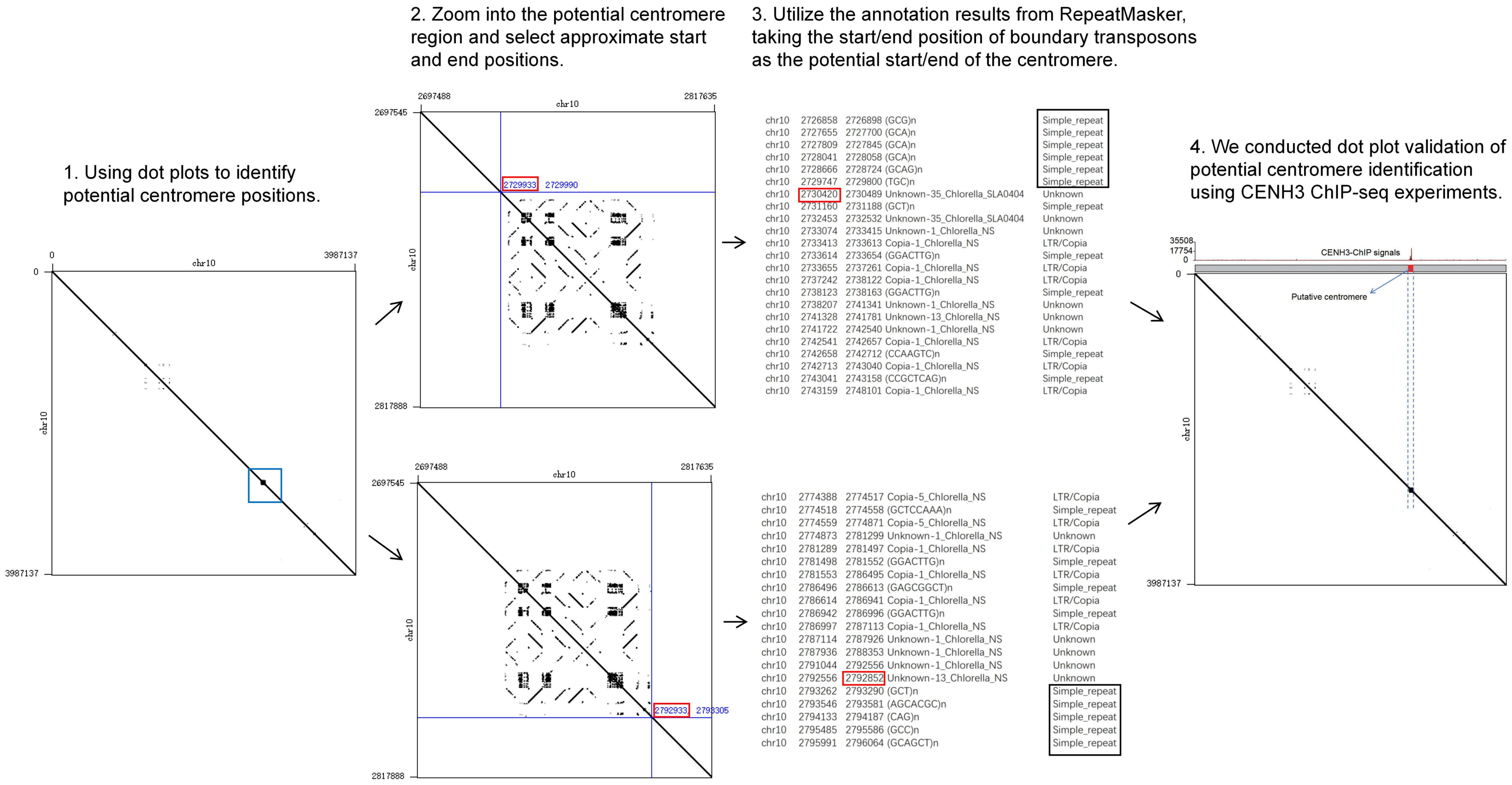


**Supplementary Fig. S10.** Overview of the dot plot method of identifying potential centromere positions in the genome, illustrated using chromosome 10 of *Chlorella sorokiniana* NS4-2.

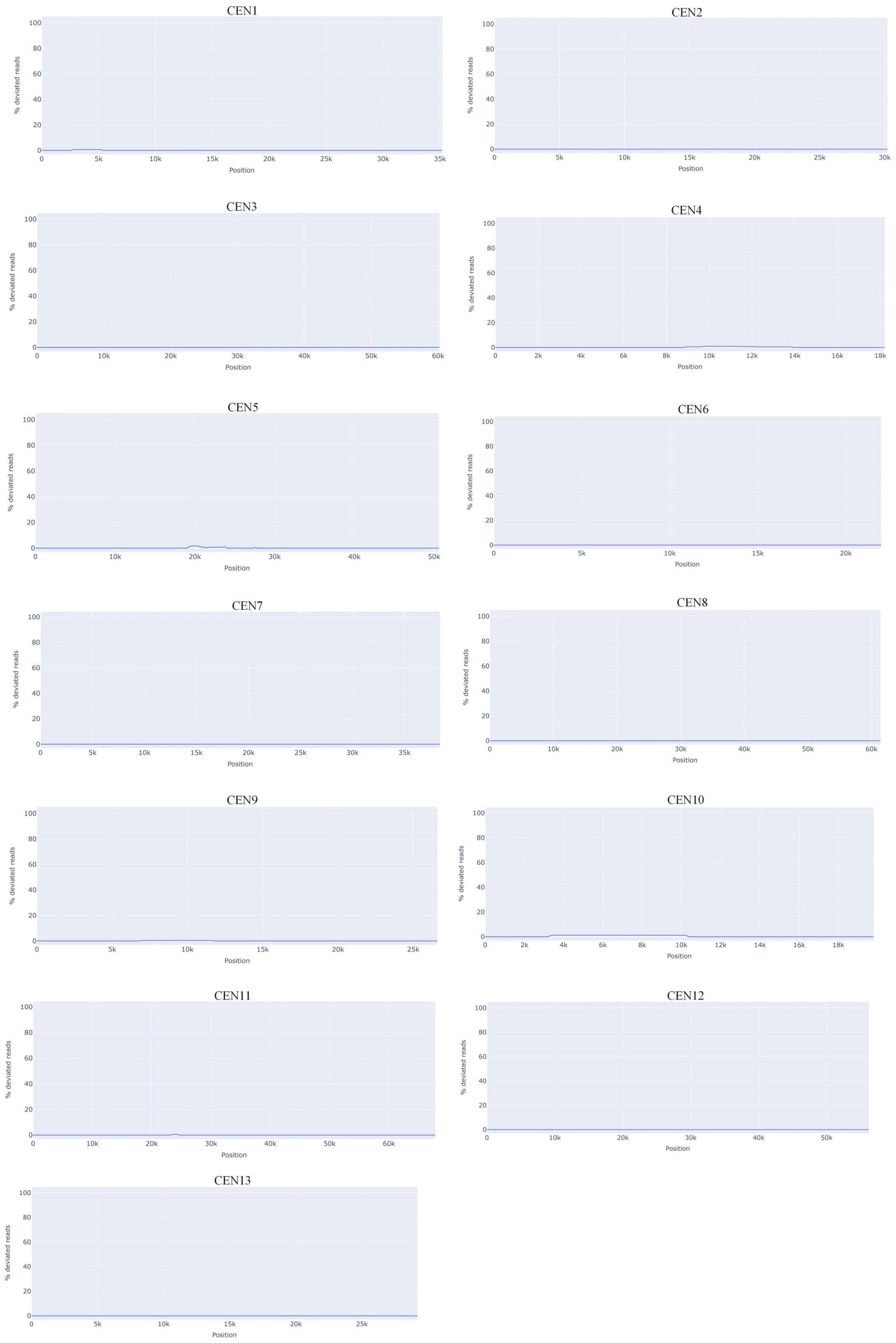


**Supplementary Fig. S11.** Evaluation of the 13 putative centromere assemblies of *Chlorella pyrenoidosa* DBH using VerityMap. The x-axis represents the position of the centromere, while the y-axis indicates the percentage of deviated reads (negligible in all cases).

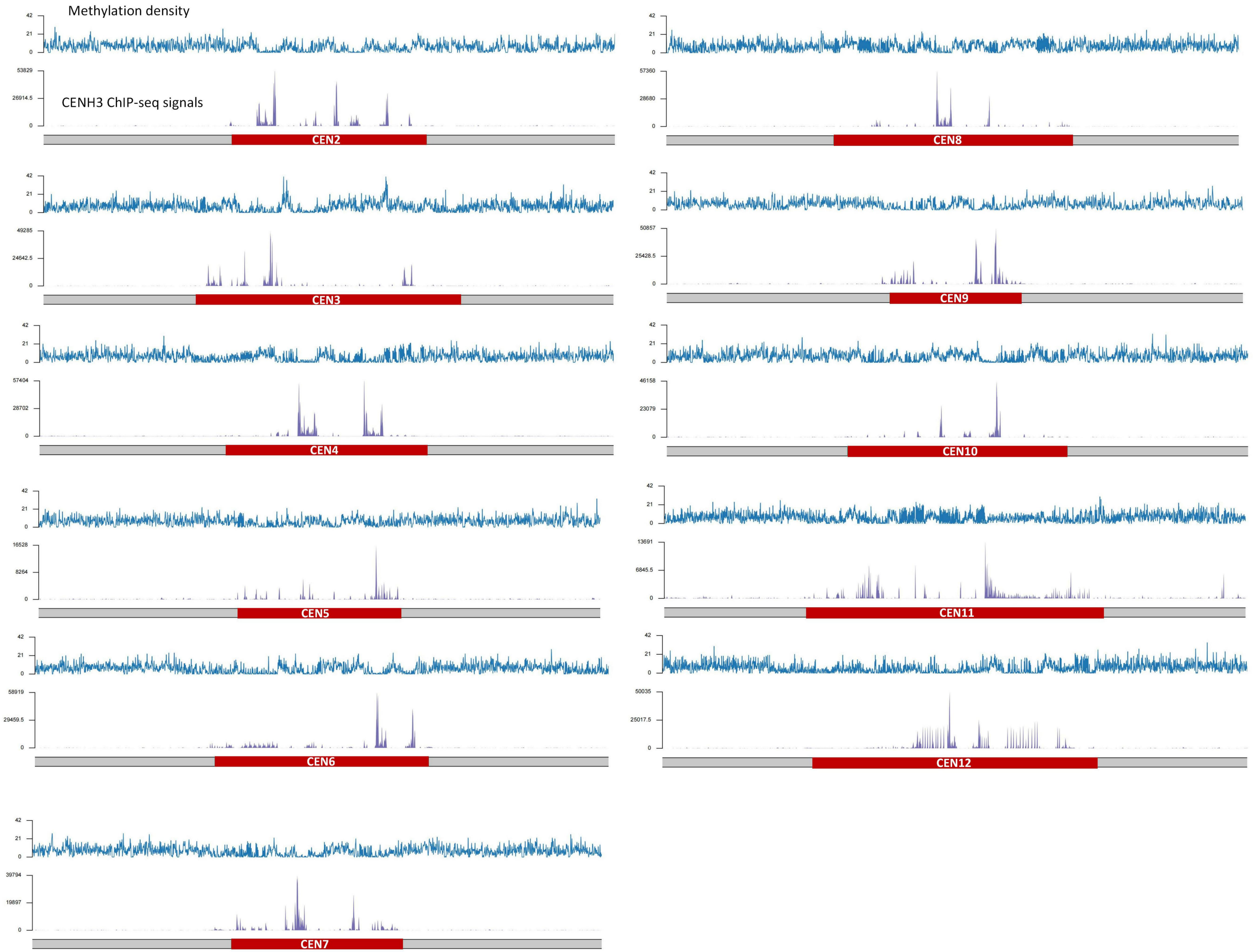


**Supplementary Fig. S12.** Centromeric methylation features of *Chlorella sorokiniana* NS4-2. Schematic overview depicting each centromere along with its 50 Kb upstream and downstream regions of *C. sorokiniana* NS4-2. The illustration shows the distribution of methylation density (determined by HiFi data), CENH3 ChIP-seq signals, and the location of centromeric regions (red bands).

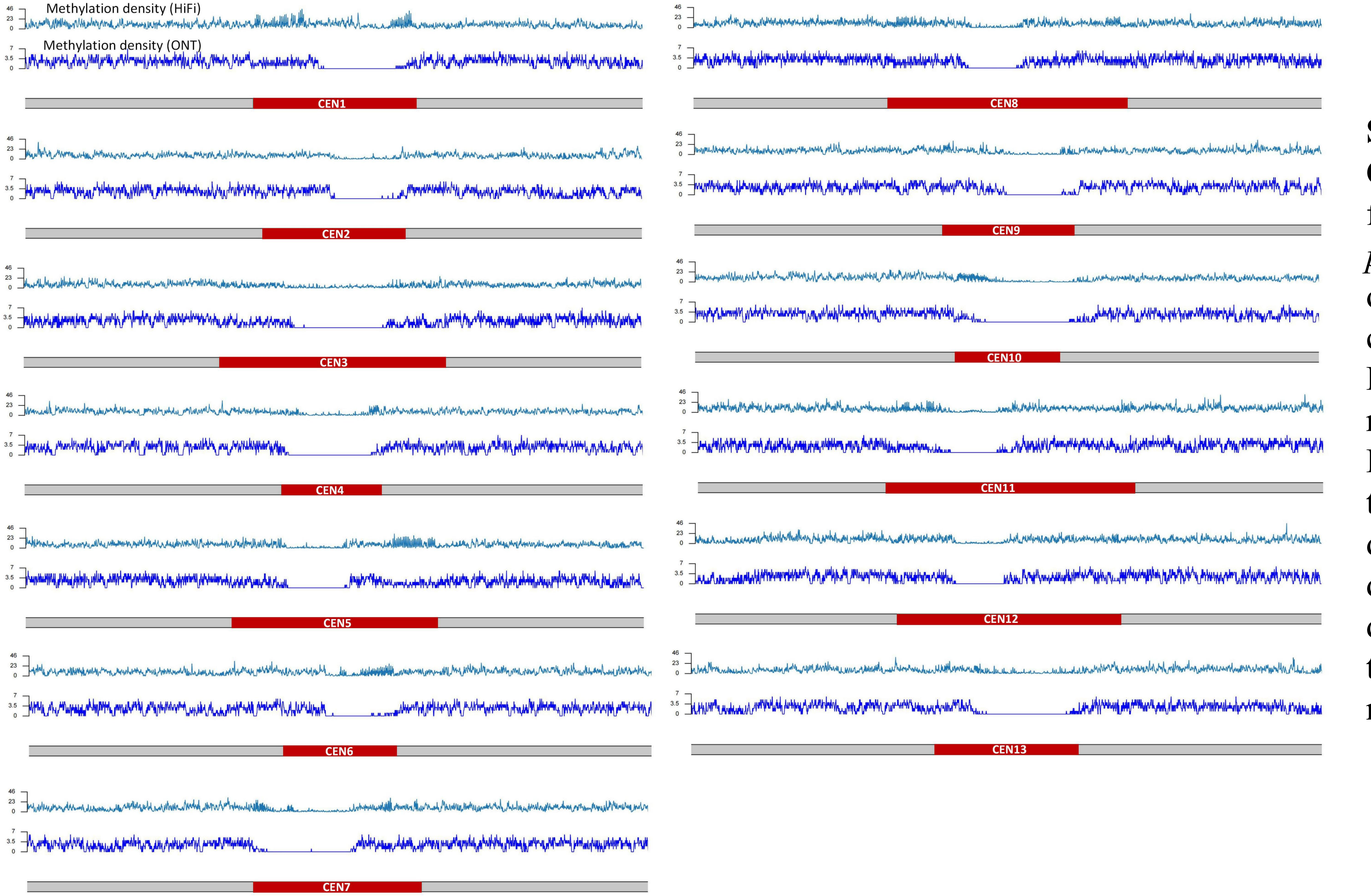


**Supplementary Fig. S13.** Centromeric methylation features of *Chlorella pyrenoidosa* DBH. Schematic overview depicting each centromere along with its 50 Kb upstream and downstream regions of *C. pyrenoidosa* DBH. The illustration shows the distribution of methylation density determined by HiFi data, methylation density determined by ONT data, and the loction of centromeric regions (red bands).

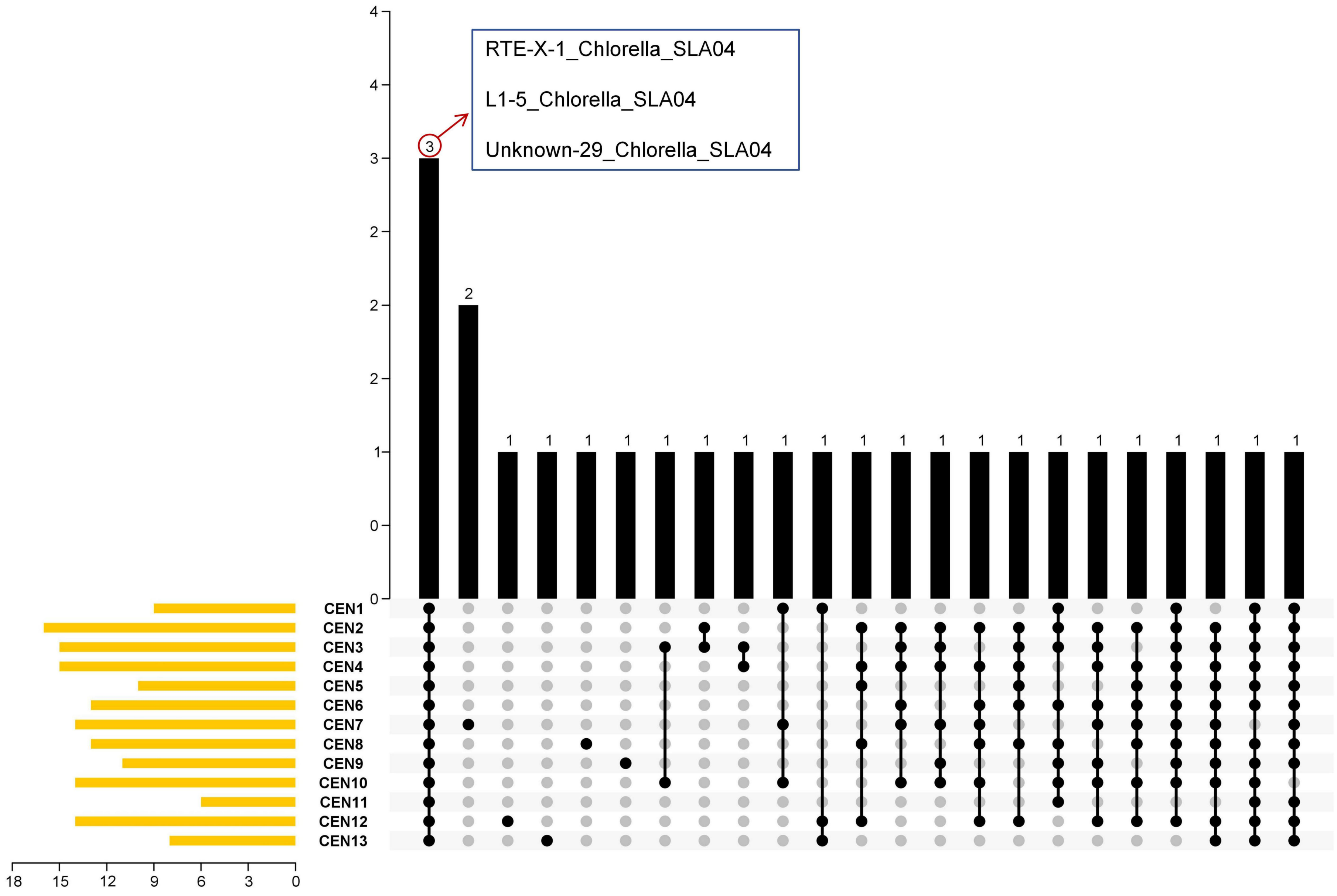


**Supplementary Fig. S14**. Upset plot depicting the overlap of centromeric repeat subfamilies among 13 centromeres (CENs) in *Chlorella* sp. SLA-04.

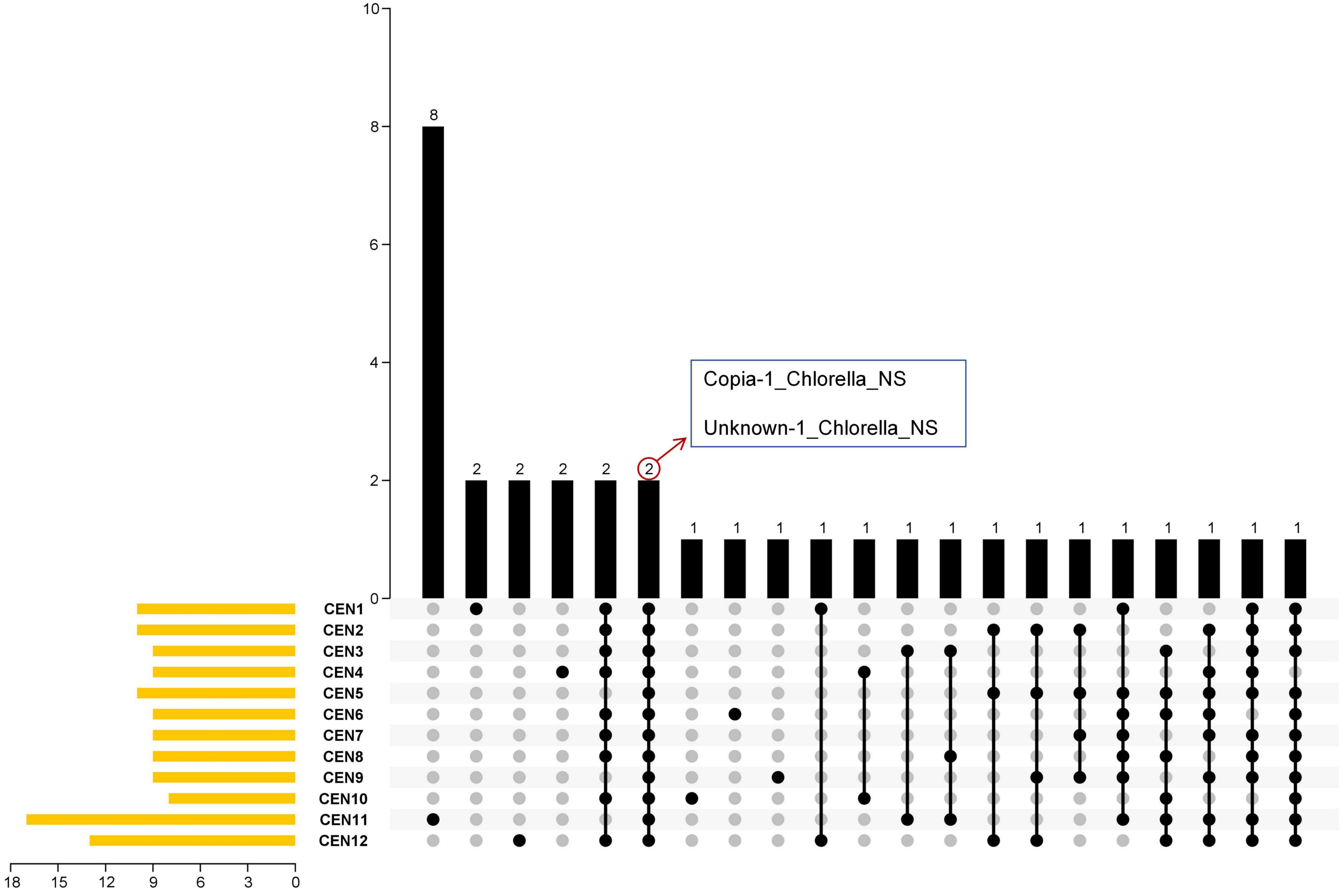


**Supplementary Fig. S15**. Upset plot depicting the overlap of centromeric repeat subfamilies among 12 centromeres (CENs) in *Chlorella sorokiniana* NS4-2.

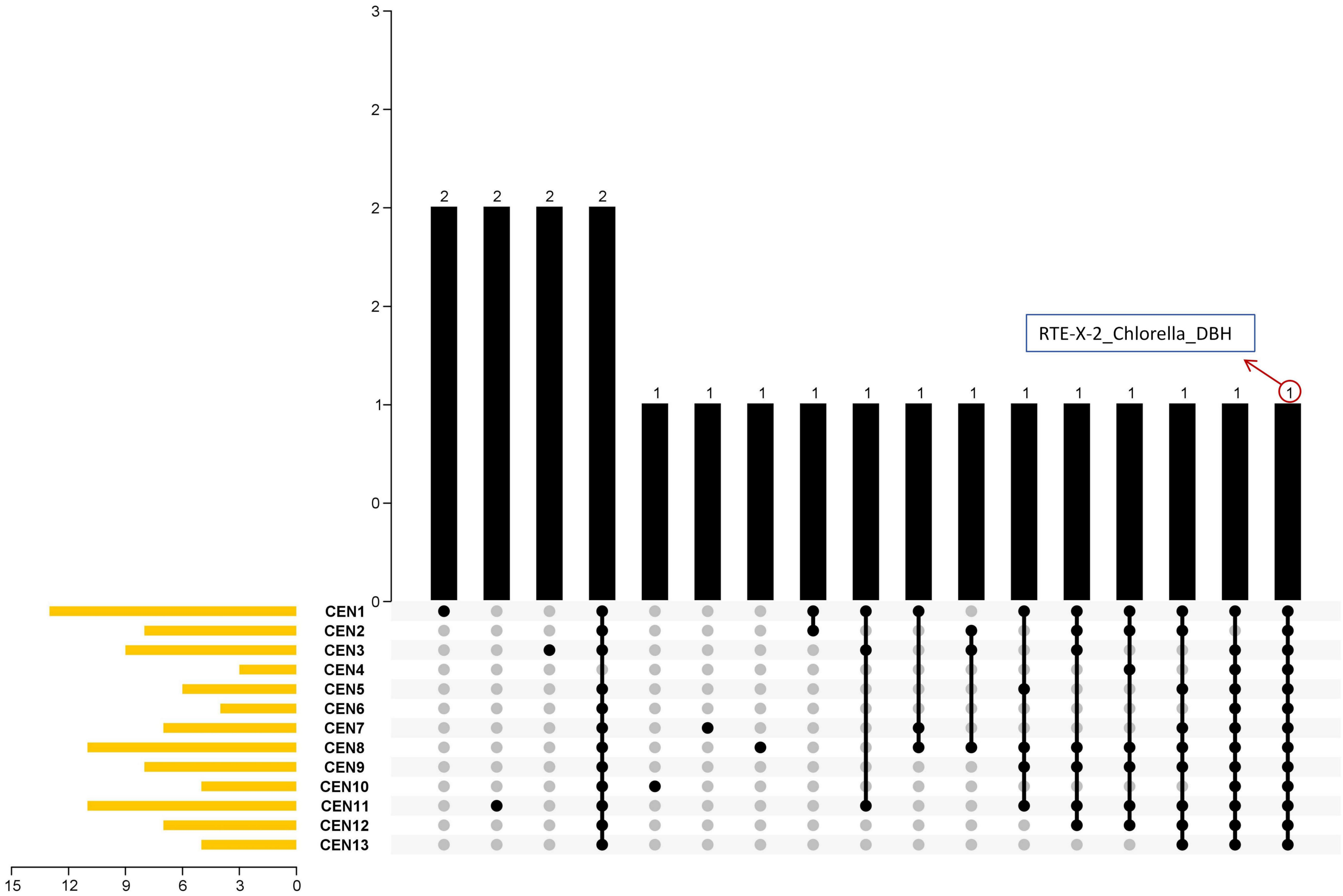


**Supplementary Fig. S16**. Upset plot depicting the overlap of centromeric repeat subfamilies among 13 centromeres (CENs) in *Chlorella pyrenoidosa* DBH.

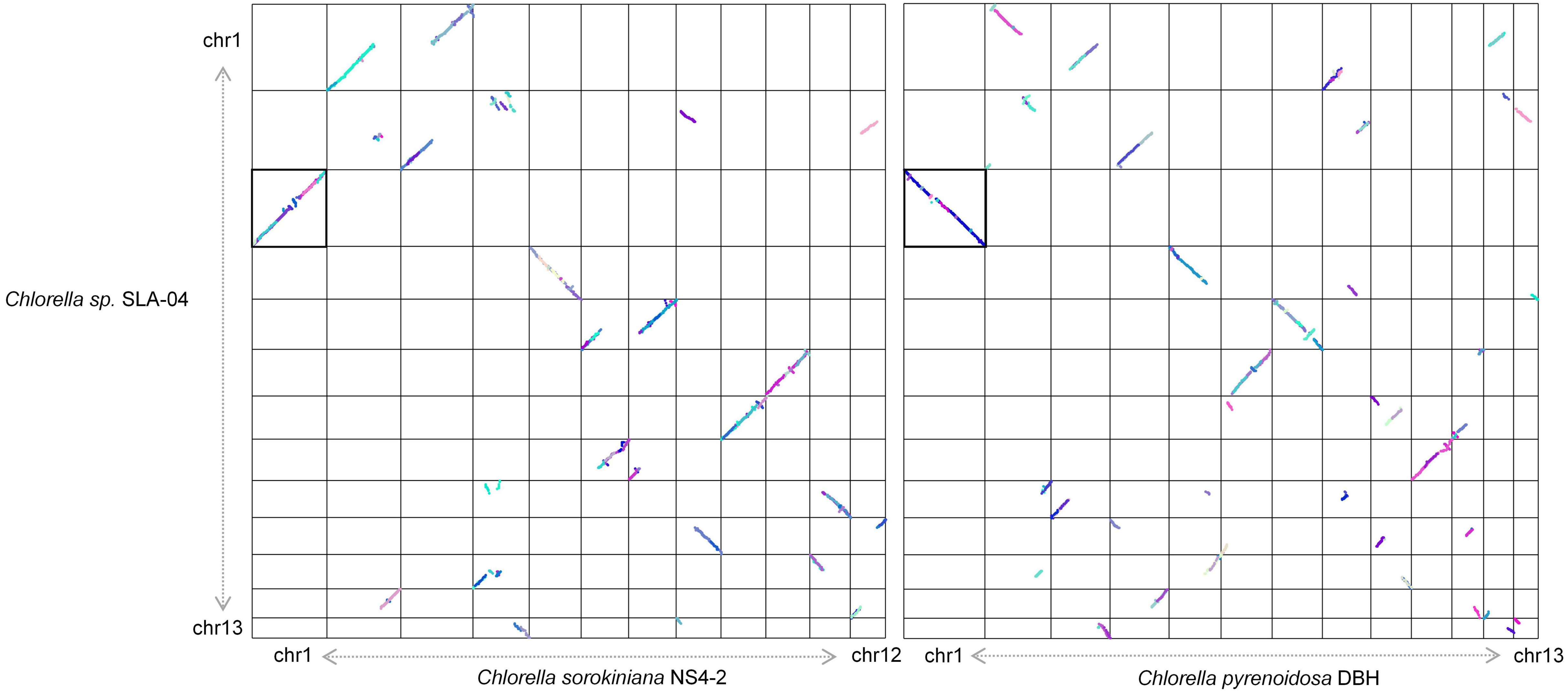


**Supplementary Fig. S17**. Dot plots of interspecies syntenic blocks of *Chlorella* sp. SLA-04, *Chlorella sorokiniana* NS4-2 and *Chlorella pyrenoidosa* DBH. Each dot indicates a syntenic gene pair detected by MCScanX. Synteny blocks were colored by MCScanX.
